# Molecular characterization of the archaic HLA-B*73:01 allele reveals presentation of a unique peptidome and skewed engagement by KIR2DL2

**DOI:** 10.1101/2024.11.25.625330

**Authors:** Philipp Ross, Hugo G. Hilton, Jane Lodwick, Tomasz Slezak, Lisbeth A. Guethlein, Curtis P. McMurtrey, Alex S. Han, Morten Nielsen, Daniel Yong, Charles L. Dulberger, Kristof T. Nolan, Sobhan Roy, Caitlin D. Castro, William H. Hildebrand, Minglei Zhao, Anthony Kossiakoff, Peter Parham, Erin J. Adams

**Affiliations:** Department of Biochemistry and Molecular Biology, The University of Chicago, Chicago, Illinois; Committee on Genetics, Genomics and Systems Biology, University of Chicago, USA; Department of Structural Biology, School of Medicine, Stanford University, Stanford, USA; Department of Microbiology & Immunology, School of Medicine, Stanford University, Stanford, USA; Department of Microbiology & Immunology, University of Oklahoma Health Sciences Center, Oklahoma City, Oklahoma, USA; Department of Health Technology, Technical University of Denmark, Kgs. Lyngby 2800, Denmark

**Author notes:** These authors contributed equally to this manuscript (Philipp Ross, Hugo G. Hilton). Authors to whom correspondence should be addressed Address correspondence and reprint requests to: Dr. Erin J. Adams, Department of Biochemistry and Molecular Biology, University of Chicago, GCIS W236, 929 E. 57^th^St. Chicago IL 60649. Email address Dr. Peter Parham, Department of Structural Biology and Microbiology & Immunology, Stanford University, Fairchild D-157, 299 Campus Drive West, Stanford, CA 94305.

**Keywords:** Natural Killer Cells, MHC, Comparative Immunology/Evolution, Antigens/Peptides/Epitopes Running title: HLA-B*73:01 allele peptidome and engagement by KIR2DL2

## Abstract

HLA class I alleles of archaic origin may have been retained in modern humans because they provide immunity against diseases to which archaic humans had evolved resistance. According to this model, archaic introgressed alleles were somehow distinct from those that evolved in African populations. Here we show that HLA-B*73:01, a rare allotype with putative archaic origins, has a relatively rare peptide binding motif with an unusually long-tailed peptide length distribution. We also find that HLA-B*73:01 combines a restricted and unique peptidome with high-cell surface expression, characteristics that make it well-suited to combat one or a number of closely-related pathogens. Furthermore, a crystal structure of HLA-B*73:01 in complex with KIR2DL2 highlights differences from previously solved structures with HLA-C molecules. These molecular characteristics distinguish HLA-B*73:01 from other HLA class I alleles previously investigated and may have provided early modern human migrants that inherited this allele with a selective advantage as they colonized Europe and Asia.

## Introduction

When modern humans migrated out of Africa they hybridized with archaic humans, including Neanderthals and Denisovans, that were already resident in Europe and Asia (1-4). As a result, 1.5-6% of modern non-African human genomes derive from archaic humans. In general, purifying selection has acted to remove archaic human DNA from modern humans (5). However, a limited number of introgressed archaic human genes have been preferentially retained (6-8). Amongst these adaptive loci are several dedicated to immune function, including selected alleles of the highly polymorphic *HLA class I* genes (*HLA-A*, *-B* and *-C*) (6). HLA class I molecules are present on the surface of all nucleated cells and are one of two primary classes of molecules encoded by the Major Histocompatibility Complex (MHC), the other being HLA class II. Their function is to display peptide fragments of varying length from within the cell for outward immunosurveillance by T cells and natural killer (NK) cells, allowing the immune system to differentiate healthy from diseased tissue. The kinds of peptides bound by an HLA class I molecule is dependent on the amino acid sequence of its peptide binding groove and underlies the correlation of specific HLA class I alleles with protection or susceptibility to a range of autoimmune and infectious diseases.

Admixture with archaic humans is hypothesized to have restored HLA diversity in modern humans following the population bottleneck that occurred during the out of Africa migration and may have been a route to acquire advantageous HLA variants already adapted to local pathogens (6, 9). A putative archaic HLA class I allele of particular interest is *HLA-B*73:01* (B*73:01). This exceptional allele, the only member of a deeply divergent lineage (MHC-BII), is distinct from the MHC-BI lineage to which all other human HLA-B alleles belong and is more closely related to subsets of chimpanzee and gorilla MHC-B than other human HLA-B alleles (6, 10). Further distinguishing B*73:01 is that it is one of only two of the HLA-B classes that encodes the C1 Natural Killer (NK) cell-binding epitope formed by residues V76 and N80 in the α1 helix (11, 12). Of note, this C1 motif is only found in B*73 and B*46, and is found in all known allotypes of these genes (4, and 100 allotypes, respectively, out of 5,745 total B proteins at last count) (13). This epitope confers reactivity with killer-cell immunoglobulin-like receptors (KIR) 2DL3 and 2DL2, inhibitory receptors that regulate the function of human NK cells. HLA-B*46:01 (B*46:01), the other HLA-B allele with a C1 epitope, acquired this epitope more recently through a mini-gene conversion of B*15 with HLA-C*01:02 (14, 15) (C*01:02). Both B*73:01 and B*46:01 have been shown to bind KIR2DL2 and KIR2DL3 using HLA-coupled beads and KIR-Fc fusion reagents (11, 16), and HLA-B*46:01 expressing cells are able to functionally inhibit KIR2DL3 expressing NK cells (14).

Carriers of the C1 epitope as well as KIR2DL2 or KIR2DL3 have varying susceptibilities to certain diseases (17). One of the earliest identified examples of this was the finding that individuals that are homozygous for the C1 epitope and KIR2DL3 are better protected from chronic HCV than individuals without C1-epitope containing HLAs or C1 carriers that also encode KIR2DL2 (18). These examples are thought to be mediated by Natural Killer (NK) cells, but KIR2DL2 is also known to be associated with adaptive immune outcomes (19). More recently, an in depth analysis of B*46:01 carriers in Southeast Asia showed that HIV + carriers of B*46:01 more rapidly progressed to AIDs than individuals without (20) and that the NK cell phenotypes of these individuals showed unusual levels of activation. These associations raise the possibility that alleles carrying the C1 epitope for KIR2DL2 or KIR2DL3 play a direct role in immunosurveillance. While there has been extensive work in understanding the interaction of these KIRs with HLA-C alleles bearing the C1 epitope, relatively little is known about how this C1 epitope in B*46:01 and B*73:01 is engaged by KIRs. Furthermore, little is known about the peptide repertoire presented by B*73:01, or how it might be associated with human disease.

Although B*73:01 presumably conferred some selective advantage when it was initially introduced into modern humans by adaptive introgression, the allele is now rare, indicating either that the selective pressure that drove its retention is no longer present, or that that selective advantage also comes with a cost. However, due to the low frequency of B*73:01 in modern day human populations (6), confident genetic associations with maladies such as infectious diseases would be underpowered and therefore unreliable. Thus, to better understand why B*73:01 was initially retained in modern humans, we studied its peptide repertoire, cell-surface expression, and three-dimensional atomic structure alone and in complex with KIR2DL2 and compared these characteristics to other HLA alleles. Our findings indicate that B*73:01 encodes a protein with a highly distinctive peptide repertoire with unique peptide presentation abilities, and is engaged by KIR2DL2 with a skewed footprint, binding that is likely modulated by the length of the presented peptide. Together, these features may have important immune consequences, which underlie its adaptive introgression into modern human genomes (6, 15, 21-23).

## Results

### B*73:01 presents a restricted peptide binding repertoire with an unusual length distribution

To investigate the repertoire of peptides bound by B*73:01, we purified heterogeneously loaded protein, eluted the bound peptides, and determined their sequences by LC-MS (24-26). We then compared B*73:01 peptides with those of other well characterized alleles: HLA-B*15:01 (B*15:01), the partner of the gene conversion that formed B*46:01, as well as HLA-C*01:02 (C*01:02), and HLA-B*46:01 (B*46:01) that were all expressed, purified, and sequenced the same way in a previous study (15). Of note, since a pan class-I antibody (*α*-W6/32) column was used for purification we do not anticipate that this purification would introduce biases to the eluted peptides compared to other HLA classes. With only 573 distinct peptides captured and confidently identified, B*73:01 has the smallest of the four peptide repertoires and is an order of magnitude smaller than the >6000 peptides bound by B*15:01 (**Figure 1A**). Furthermore, fewer than 10% of the B*73:01 bound peptides overlapped with one or more of the other alleles investigated. Examples of peptides eluted from B*73:01 and used later in this analysis are listed in **Table S1**.

**Figure 1.**
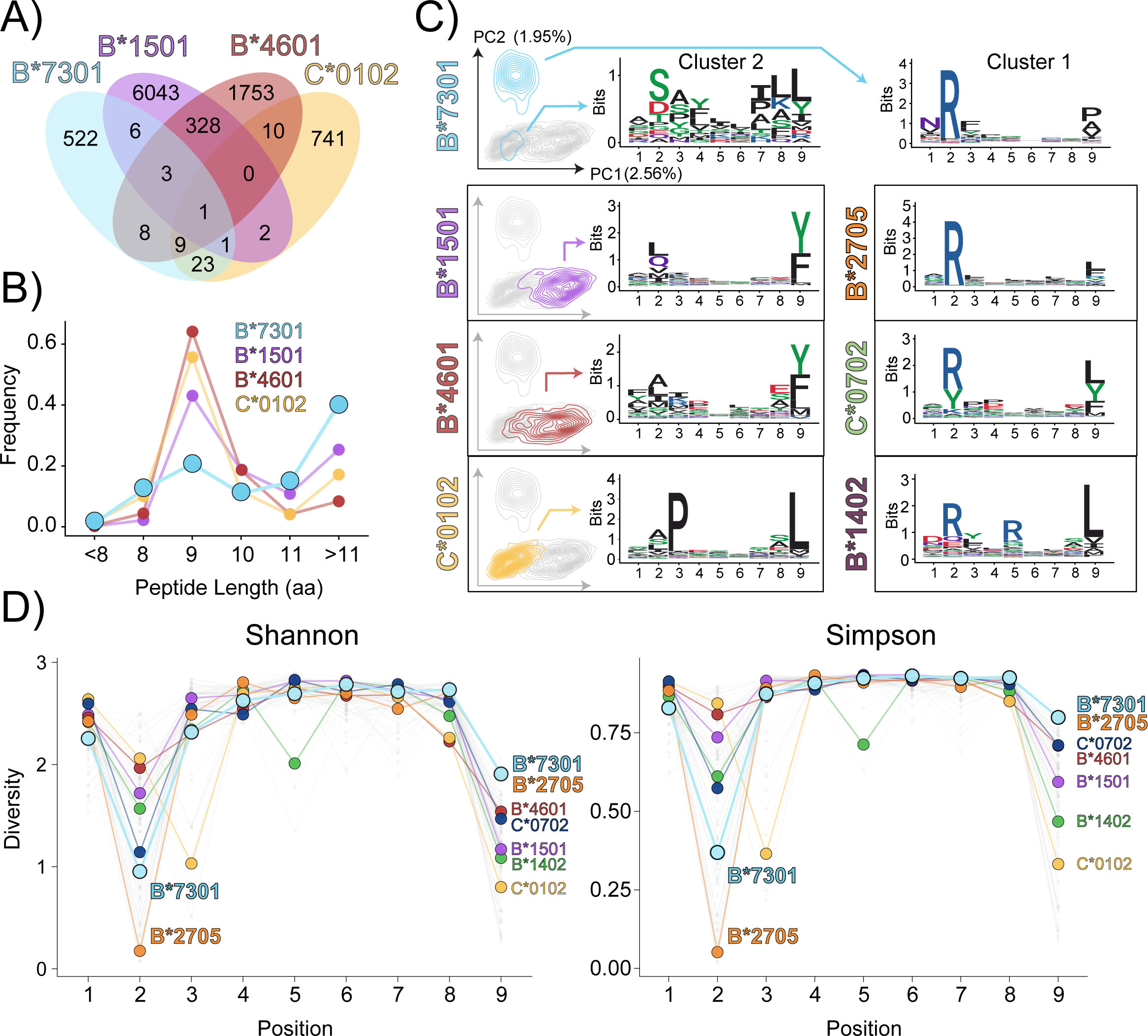
HLA-B*73:01 has evolved a rare peptide binding repertoire with an unusual length distribution. **A)** Venn diagram showing the number of distinct peptides eluted from the HLA class I allotypes: B*73:01, B*15:01, B*46:01 and C*01:02. The number of peptides shared with each of the other allotypes is shown. **B)** Length distribution of peptides eluted from the HLA allotypes in **(A)**. **C)** Principal components analysis contour plots show the distribution of nonamer peptides bound by each HLA class I allotype (on left). For comparison, in each plot the distribution of nonamer peptides from the other allotypes is shown in grey. The percentage of the variation in the dataset shown by the first (PC1) and second (PC2) principal components is shown near the axises. Sequence logos of 9mers isolated from the analysis in **(A)**, categorized by unsupervised alignment and clustering, are shown for peptides eluted from HLA-B*73:01, B*15:01, B*46:01 and C*01:02. Data from Sarkizova et al. (32) was used to generate similar plots for B*27:05, C*07:02 and B*14:02 (on right). The height of each amino acid in the plot represents the level of conservation at that position and its relative frequency. The colors of each amino acid correspond to their biochemical characteristics: acidic (red), basic (blue), hydrophobic (black) and polar (green). **D)** Shannon and Simpson diversity indices calculated for each position for HLA-B*73:01, B*15:01, B*46:01 and C*01:02 using peptides eluted and sequenced by mass spectrometry in our study.

Contributing to the distinctiveness of B*73:01 peptides is a considerable promiscuity in the eluted peptide length. When compared to other alleles, we found that nonamer peptides comprise only ∼20% of B*73:01 peptides (**Figure 1B**). Instead, the majority (∼40%) of peptides that B*73:01 presents are greater than 11 amino acids in length, reminiscent of the length of peptides that are bound to HLA class II molecules, which, by virtue of their open-ended peptide binding groove, routinely bind longer peptides (27). To understand how B*73:01 can accommodate such long peptides, we compared the peptide binding motifs of peptides of varying lengths and found that even as the peptides get longer, they retain a canonical B*73:01 binding motif with a preferred arginine at Position 2 (P2) and a hydrophobic residue at the peptide C-terminus (**Figure S1A**). This suggests that these positions remain the primary peptide anchors in B*73:01 with the intervening residues accommodated by “bulging out” from the peptide binding pocket without greatly destabilizing the protein complex. Peptides of lengths above 11 amino acids were analyzed for strong HLA-B73:01 binding using NetMHCpan-4.1 (28). Several of these longer peptides (∼12-15 amino acids long) were predicted to exhibit high binding values (**Table S1**). Because nonamers represent the most common length of peptide from classical HLA I class I proteins and the nonamer sequences from B*73:01 maintain the consensus motifs represented in the other lengths of B*73:01 peptides, we chose to compare this length with those of other nonamers from other HLA alleles in the analysis presented below.

Visualization of the relative similarity of the nonamer peptidomes of B*73:01, B*15:01, B*46:01 and C*01:02 by principal components analysis supports the uniqueness of the B*73:01 peptide repertoire motifs, showing that most B*73:01 bound peptides have little spatial overlap with those of the other three alleles (**Figure 1C; left column**). The greatest contributor to PC2 is the hydropathy index (55%), a measure of the hydrophobicity/hydrophilicity of side chains; in B*73:01 this is reflected in the strong preference for an arginine at P2 (**Figure 1C**). Using an unsupervised alignment and clustering of peptide sequences (29, 30), we can identify 2 divergent peptide binding motifs for B*73:01, with 85% of nonamers contributing to Cluster 1 and the remaining peptides to Cluster 2. The Cluster 1 peptides maintain this P2-Arginine/P9-hydrophobic preference, while peptides from Cluster 2 show less specific anchor preferences yet are more similar to peptides bound by B*15:01, B*46:01, and C*01:02. The motif represented in the Cluster 1 nonamer peptides from B*73:01 shows strong resemblance to peptides bound by the HLA-B*27 family of alleles (for example B*27:05, shown in orange in **Figure 1C**, right panel). Of note, there is evidence that other HLA alleles with compact peptidomes, like B*27:05, although sometimes protective, may also predispose individuals to autoimmune disease, as is the case with B*27:05 and axial spondyloarthritis (31).

To understand how B*73:01 compares to HLA alleles representative of those with varying frequencies worldwide, we compared our B*73:01 peptidome data to that from Sarkizova et al., which includes >185,000 total peptides from 95 alleles, chosen such that 95% of individuals worldwide have at least one of the HLA-A, HLA-B and HLA-C alleles included (32). The peptidome for each allele was generated using the same parental cell line (BLCL 721.221) as we used for B*73:01, B*15:01, B*46:01, and C*01:02. To confirm that we can confidently combine these two datasets together we compared the peptide binding motifs of eluted nonamers for alleles in both datasets. By doing this, we found that eluted nonamers from both studies were highly concordant with Pearson correlation values of 0.93, 0.93, and 0.99 for B*15:01, B*46:01, and C*01:02, respectively (**Figure S1B**), suggesting that the peptidome of B*73:01 sequenced using our approach is comparable to those of Sarkizova (33). Incorporating these additional peptide repertoires into our analysis, the similarity between the peptide preferences of B*73:01 with those of other HLA alleles became apparent. For nonamer peptides, the HLA alleles B*27:05, C*07:02 and B*14:02 share commonalities in their preference for an arginine at Position 2 and a hydrophobic residue (leucine) at Position 9 (**Figure 1C; right panel**).

Using a similar approach to Sarkizova et al. we compared B*73:01 to this more extensive peptide repertoire analysis using the data from both peptides bound (“peptide space”) and the variable pocket residues within the platform regions of HLA molecules (“pocket space”). These analyses revealed at a more granular level which peptidomes shared the greatest similarity to that of B*73:01. Consistent with our previous findings in **Figure 1**, HLA B*27:05 (B*27:05) had the highest similarity to B*73:01 in both peptide space and pocket space (**Figure S2A**), whereas HLA-C*07:02 (C*07:02), HLA-C*07:01 (C*07:01) and HLA-C*06:02 (C*06:02) had high similarity in peptide space (**Figure S2A**; left panel), and HLA-B*56:01 (B*56:01) and HLA-B*14:02 (B*14:02) had highest similarity in terms of pocket space. B*07:02, B*07:04, B*08:01, B*13:02, B*40:02, B*40:06, B*42:01, B*54:01, B*55:01, B*55:02, and B*56:01 also shared in similarity to B*73:01 in terms of pocket space (labels not shown in figure due to space constraints) (**Figure S2A; right panel**). Thus, similar to comparisons made with only B*15:01, B*46:01, and C*01:02, the divergent nature of the B*73:01 peptidome stems from its strong preference for arginine at P2. Re-analysis of the peptide repertoire length of B*73:01 with this expanded dataset recapitulated our original results, notably a preference for peptides longer than 11 amino acids is a unique feature of B*73:01 (**Figure S2B**).

Recent reports suggest that HLAs can be categorized as either ‘generalists’ or ‘specialists’ depending on the promiscuity of the peptides they can present (34). To compare the diversity of these B*73:01 characterized peptides we calculated diversity indices using either Shannon (35) or Simpson measures at each position of the HLA alleles presented above and those from Sarkizova et al. (**Figure 1D**). Interestingly, while these analyses show that B*73:01 has relatively low sequence diversity at P2 (with B*27:05 having the lowest diversity), it and B*27:05 have the highest sequence diversity of all these alleles at position 9. Together, these data show that B*73:01 has evolved the ability to present peptides with a restricted binding motif relative to other more common alleles, similar to that seen for the B*27 family of alleles. Additionally, B*73:01 peptides are proportionally longer and vary most at their C-terminal anchor, while being heavily biased towards arginine at the P2 anchor position.

### B*73:01 combines a size-restricted and unique peptidome with above average cell-surface expression

Recent work suggests that peptide binding promiscuity of a given HLA class I peptidome is inversely correlated with its cell surface expression (36). Considering the restricted repertoire of the peptides presented by B*73:01, we hypothesized that it would have concordantly high cell-surface expression levels relative to other alleles. To test this, we first investigated the expression of each of the alleles used in our original analysis (**Figure 1A-C**) (15) in 721.221 cells (which lack endogenous expression of HLA class I) using W6/32 (37), an antibody that recognizes all HLA class I alleles with equal avidity. Indeed, we find that B*73:01 is the highest expressed allele of the four, being expressed at a significantly higher level than both C*01:02 and B*46:01, but at an approximately equal level to B*15:01 (**Figure 2A**). The low cell-surface expression of C*01:02 is consistent with previous studies showing that alleles of the HLA-C locus are expressed at a significantly lower level at the cell-surface than alleles of the HLA-A or HLA-B loci (38, 39). Contributing to this effect is the KYRV motif (**Figure 2B**; highlighted), a four amino acid motif at positions 66, 67, 69 and 76 in the α1 helix that is present in all HLA-C, but not in HLA-A or HLA-B (40). Because the gene conversion that created B*46:01 inserted the KYRV motif into the backbone of B*46:01, we hypothesized that this may be the source of the difference in expression between B*73:01 and B*46:01. To test this, we examined the expression of swap mutants in HeLa cells in which the residues at positions 66, 67 and 69 (both alleles encode Val76) present in B*46:01 were replaced with those found in highly expressed B*73:01 (which lacks the KYR motif and encodes ICA at these residues; **Figure 2B**) and compared them with their wildtype counterparts. Consistent with the results from 721.221 cells, we found that in HeLa cells, wild-type B*73:01 is expressed at a significantly higher level than B*46:01 (**Figure 2C**). Further, we found that the B*73:01-KYR mutant had significantly reduced expression and the B*46:01-ICA mutant had significantly increased expression compared to their respective wild-types (**Figure 2C**). These results are consistent with the KYRV motif playing a significant role in determining the divergent levels of cell-surface expression between these two C1+ HLA-B alleles. Thus, while B*73:01 and B*46:01 both encode the C1 epitope and serve as ligands for KIR2DL3, they differ considerably in their α1 domain sequence, which not only correlates with their distinctive peptidomes, but also with their divergent cell-surface expression.

**Figure 2.**
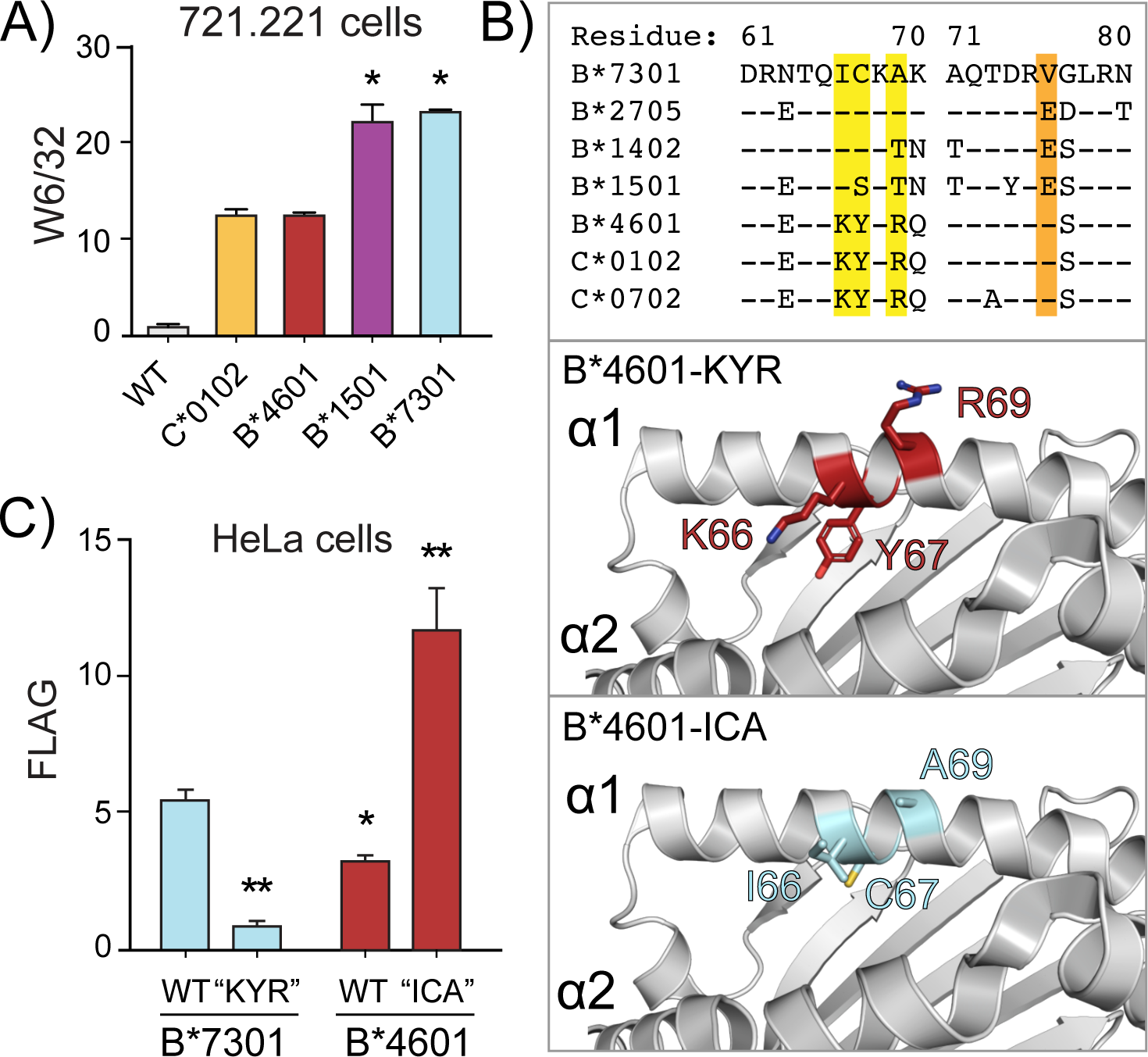
HLA-B*73:01 combines a size-restricted and unique peptidome with above-average cell-surface expression. **A)** Cell-surface expression of 4 HLA class I allotypes as determined by W6/32 staining in transfected 721.221 cells (asterisks indicate expression that differs significantly from HLA-B*46:01 (two-tailed t-test, * indicates p<0.005). **B) Top panel:** An alignment of selected HLA amino acid sequences from residue 61-80, containing the KYRV motif (highlighted in yellow and orange) known to affect the cell surface expression of HLA-C alleles (ref). A crystal structure of HLA-B*46:01(PDB ID: 4LCY) showing the wildtype KYR motif in red (**middle panel**) or the ICA motif in cyan (**lower panel**), following *in silico* mutagenesis in PyMol. **C)** Surface expression of flag-tagged wild-type (WT) HLA-B*73:01 and HLA-B*46:01 and position 66, 67 and 69 (KYR/ICA) swap mutants in HeLa cells as determined by anti-Flag Ab staining; asterisks indicate expression that differs significantly from HLA-B*73:01-WT (two-tailed t-test, * indicates p<0.05, ** indicates p<0.005).

### Three dimensional structures of B*73:01 reveal the molecular basis for peptide presentation

The exceptional number of polymorphisms between class I HLA alleles is what largely confers upon them their unique antigen presenting abilities. Amino acid diversity is most pronounced in the peptide binding groove, suggesting that different environments select for the presentation of different peptides. Since the peptidome of B*73:01 is so unique, we sought to elucidate its three-dimensional structure to determine how its molecular architecture contributes to shaping its unusual peptide repertoire. Crystals of recombinant B*73:01 alone were difficult to obtain, regardless of the synthetic peptide loaded. Thus, two alternative strategies were utilized to obtain high resolution protein structures. First, we generated high affinity synthetic antibody fragments (Fabs) to HLA-B*73:01. Using this approach, we successfully generated seven Fabs that bound to B*73:01, each with nanomolar affinity (**Table S2**). While these reagents did not help as chaperones for crystal formation, as hoped, we copurified two of them (B.1 and B.8) with recombinant B*73:01 loaded with the nonamer peptide P2R (NRFLGDYVV) (**Figure S3A**) to generate a three-dimensional map by cryogenic electron microscopy (cryoEM) to a resolution of 3.1 Å (**Figure S3B**, **Table S3**). Second, we were able to solve the crystal structure of B*73:01 in complex with KIR2DL2, loaded with a 10mer peptide, NRFAGFGIGL (KP1), to 2.9 Å resolution (**Table S4**).

Our model of B*73:01 generated using these two structures shows that it adopts an overall conformation similar to that of other class I HLA alleles (**Figure 3A**) with adequate resolution to confidently place the peptides in both structures (**Figure S4**). The backbone RMSD of the two structures of B*7301 was also quite small (0.746 Å over 274 Cα atoms), indicating that presentation of different peptides and receptor binding did not alter the three-dimensional structure of B*73:01 significantly. Aside from hydrogen-bonding (h-bonding) between N114 of B*73:01 and P7Y of the P2R peptide and a pi-stacking interaction between W147 of B*73:01 and P6F of KP1, the networks of bonds (h-bonds, VDWs and salt bridges, **Table S5**) that secure P2R and KP1 within the B*73:01 binding groove are biased towards the N-and C-terminal ends of the peptide ligand (**Figure 3B**), as is the case for most other class I HLA molecules. This bias is also consistent with trends seen in the peptide-binding motif where preferences are biased towards P2 and the C-terminal position (PΩ) of the peptide (**Figure 1C**). Unlike most other alleles, B*73:01 has a slight preference for an asparagine at P1 of the peptide (**Figure 1C**); both peptides in our B*73:01 structures have this residue in the P1 position. In these two models, the asparagine side chain forms contacts with residues from the B*73:01 α helices, engaging residues on either side of the peptide binding groove, namely N63 and E163 (**Figure 3B, 3C** left panels), whereas P1-N of P2R is further stabilized through a contact with R62 on B*73:01. Thus, our structures have revealed the molecular basis of B*73:01 peptide presentation of two peptides of differing length and demonstrate an alternative network of N-terminal stabilizing interactions that function to accommodate peptides of different sequence and length.

**Figure 3.**
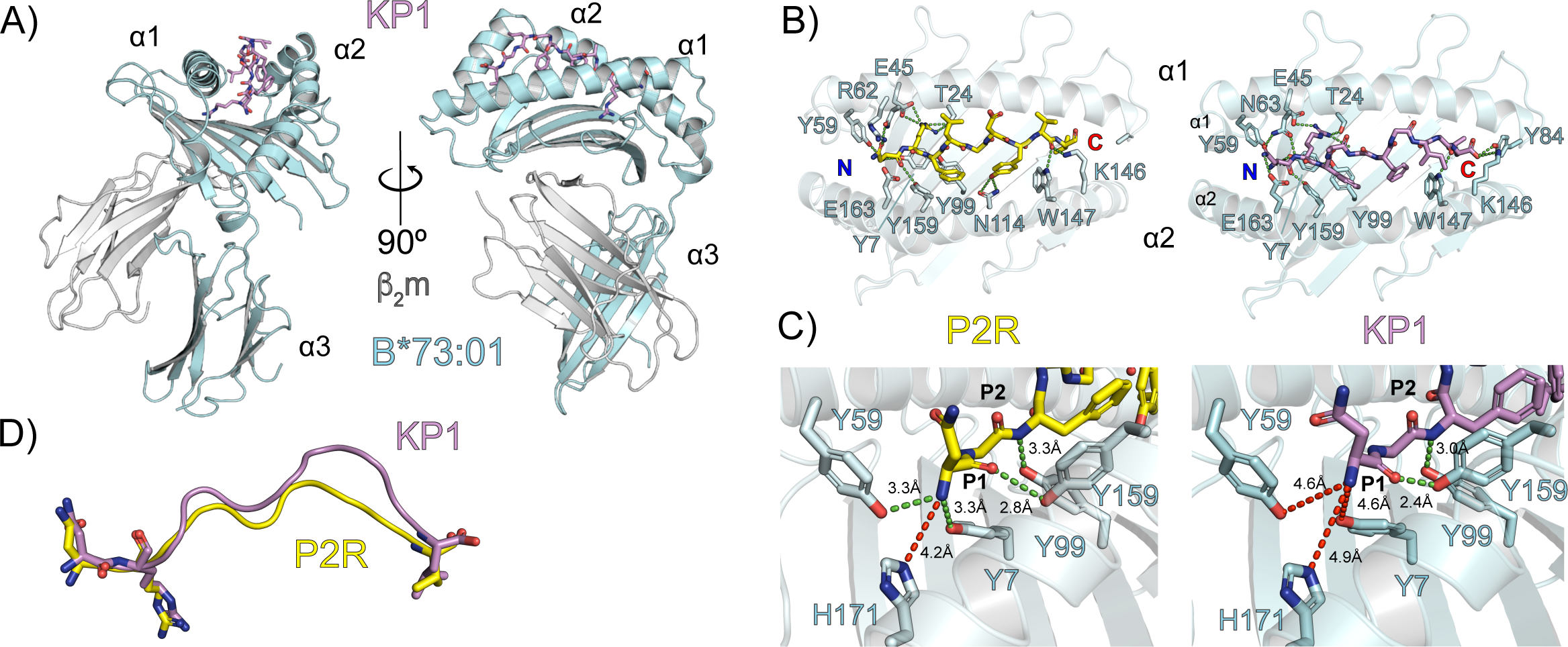
High resolution protein structures of HLA-B*73:01 bound to a 9mer and 10mer highlight flexibility within the peptide:MHC hydrogen-bonding network. **A)** Shown are two views of the structural model generated from the crystal structure of B*73:01 with the KP1 peptide. The heavy chain of B*73:01 is shown in ribbon format in cyan, β2m is shown in ribbon format in grey and the KP1 peptide is shown as sticks in mauve, with oxygens colored red and nitrogens, blue. **B)** Comparison of the platform domains of B*73:01 presenting the P2R (NRFLGDYVV) peptide, solved by cryo-EM structure (left) and that of B*73:01 with the KP1 peptide (NRFAGFGIGL). The hydrogen bonding network between B*73:01 and the P2R and KP1 peptides are indiated by dashed green lines. **C)** Atomic distances between the amino terminus of each peptide and the corresponding residues forming the A pocket with distances in angstroms. Distances in red are outside the conventional hydrogen bonding cutoff of ∼3.5Å. **D)** Comparison of the backbones of the P2R and KP1 peptides from a superposition based on the B*73:01 backbone. Peptide backbones are shown in cartoon format, colored as in **B)** & **C)**. Side-chains of the anchor residues are shown in stick format, with the coloring convention as in **A**). Shown is the displacement distance, in Angstroms, of the P5 position of the peptides.

Comparing the two structures of B*73:01 with different peptides, we can also see examples of how peptides of different sequence and length are accommodated within the B*73:01 groove. At the N and C-termini of the peptide, almost all h-bonding contacts are conserved across the two structures within our resolution limits (**Table S5)**. The only specific h-bonding contacts unique to each structure are those between the P2R peptide and Y59, R62 and N114 and the KP1 peptide and N63, N80 and Y84 (this side chain was not resolved in the P2R structure) (**Figure 3C**). The P2R peptide is a 9mer, while KP1 is a 10mer, and the difference in lengths between these two peptides are accommodated through a displacement that is biased towards the C-terminal end of the peptide. Indeed, the Ca backbone displacement of KP1 in relation to P2R at position 7 (glycine and tyrosine, respectively) is ∼4.4Å (**Figure 3D**). This tyrosine at P7 of the P2R peptide forms a unique contact with N114 of B*73:01, further helping to anchor this in the groove (**Figure 3B**).

Closer inspection of the hydrogen-bonding network between the backbone of the N-terminus of the P2R and KP1 peptides and B*73:01 revealed an adaptation that alters the canonical N-terminal anchoring observed in most classical class I structures in a length-dependent fashion (**Figure 3C**). In most class I HLA molecules, tyrosines at positions 7, 99, 159, and 171 coordinate the amine group of the N-terminus of their peptide ligands(41). In B*73:01, tyrosine 171 is replaced by a histidine; with the P2R peptide, h-bonding between the amino group and Y7 is maintained and even enhanced with Y59 (3.3Å) (**Figure 3B**). However, the N-terminus of KP1 is positioned outside the distance of relevant hydrogen-bonding for these residues (**Figure 3C**). In relation to the positioning of other canonical peptide N-termini, the N-terminus of KP1 is shifted up by an average of 1.64 Å relative to nonamers (**Figure S5A**; left panel) and an average of 1.79 Å relative to other 10mers, an 11mer, and a 15mer (**Figure S5A**; right panel). This contrasts with what is seen for the alleles B*15:01, C*01:02, and B*27:05, where each allele has both Y7 and Y171 within reasonable distance to form hydrogen bonds with their respective peptide ligands (**Figure S5B**, left panels). B*14:02, in contrast, carries H171 and shows a similar bonding network to B*73:01 as well as a similar slight preference for asparagine or aspartic acid at P1 of its peptide ligands (**Figure S5B**, right panel). Previous studies investigating peptide binding to three alleles of HLA-B*51 that carry either H171 or Y171 show that while the difference had minimal effect on peptide binding, it did have a functional effect on T cell responses (42, 43). Thus, the A-pocket of B*73:01 carries H171 instead of Y171, which likely alters the conformation of bound peptides enough to affect T cell responses, but not its peptide repertoire.

Furthermore, our analysis of peptides of varying lengths presented by B*73:01 discussed above showed conserved anchor residues near the N and C-termini suggesting that as presented peptides increase in length, they likely bulge out from the binding groove to be accommodated. In our B*73:01/KP1 structure, the conformation of the KP1 peptide (a 10mer) is displaced from the groove near the C-terminus by an average of 3.11 Å in comparison to 7 different nonamer peptides from other HLA structures (**Figure S5A**, left panel). When compared to peptides of at least 10 amino acids in length, KP1 is further displaced in this location by an average of 3.57 Å (**Figure S5A**, right panel). Together, these data suggest that the peptide binding mode of longer peptides presented by B*73:01 differs from that of most other alleles in its anchoring at the N-terminus within the A-pocket. Furthermore, B*73:01 accommodates peptides of extreme length (11+ residues) through a bulged conformation towards the C-terminal end of the peptide. While we cannot rule out that the binding of KIR2DL2 has modified the conformation of the peptide in this structure, other structural comparisons between bound and unbound HLA molecules have shown little if any peptide movement when KIR bind. For example, a comparison of crystal structures of C*07:02 loaded with the same peptide that crystallized alone, in complex with KIR2DL2, or in complex with KIR2DL3 shows minimal differences in the peptide backbone and sidechains, suggesting that binding by KIR2DL2 likely does not alter the mode of presentation of KP1 presented by B*73:01 (16) (**Figure S6**).

### The B and F pockets of B*73:01 illuminate constraints and flexibility in peptide binding

B*73:01 is part of an ancient allelic lineage of HLA-B alleles and the only member of this lineage in humans (6). The B pocket (the canonical pocket that binds the P2 of the peptide) of B*73:01 uses very similar residues as those of B*27:05 and B*14:02 (**Figure 4A**, left panel). An investigation of our structural model of B*73:01 shows that T24 and E45 are likely very important for anchoring the P2 arginine within the B pocket (44) (**Figure 4A, 4B**, left panel, positions highlighted in yellow) for both the P2R and KP1 peptides. This is also similar to what is seen in B*27:05 and B*14:02 (**Figure 4C**, **D** left panel). Notably, a previous study performing quantitative peptide binding studies tested dozens of different peptides and their ability to stabilize the B*73:01 trimeric complex in solution (25). Surprisingly, although the majority of peptides that bound most strongly contained an arginine at P2, the highest affinity binder contained a glutamic acid at P2. To reconcile this with our data, we noticed that B*73:01 also carries H9 and K70 (**Figure 4A**, left panel, positions highlighted in blue), two basic residues which are not involved in anchoring the peptides with a P2 arginine which may be recruited in stabilizing a P2 glutamic acid. Along these lines, B*40:02 is a B allele that strongly prefers a glutamic acid anchor at P2 and also carries H9 and N70, both of which form hydrogen bonds with the peptide anchor residue (**Figure 4A, S7**). Thus, B*73:01 may, in rare cases, be able to adopt its peptide binding groove to present peptides anchored at P2 with either strongly basic or acidic properties.

**Figure 4:**
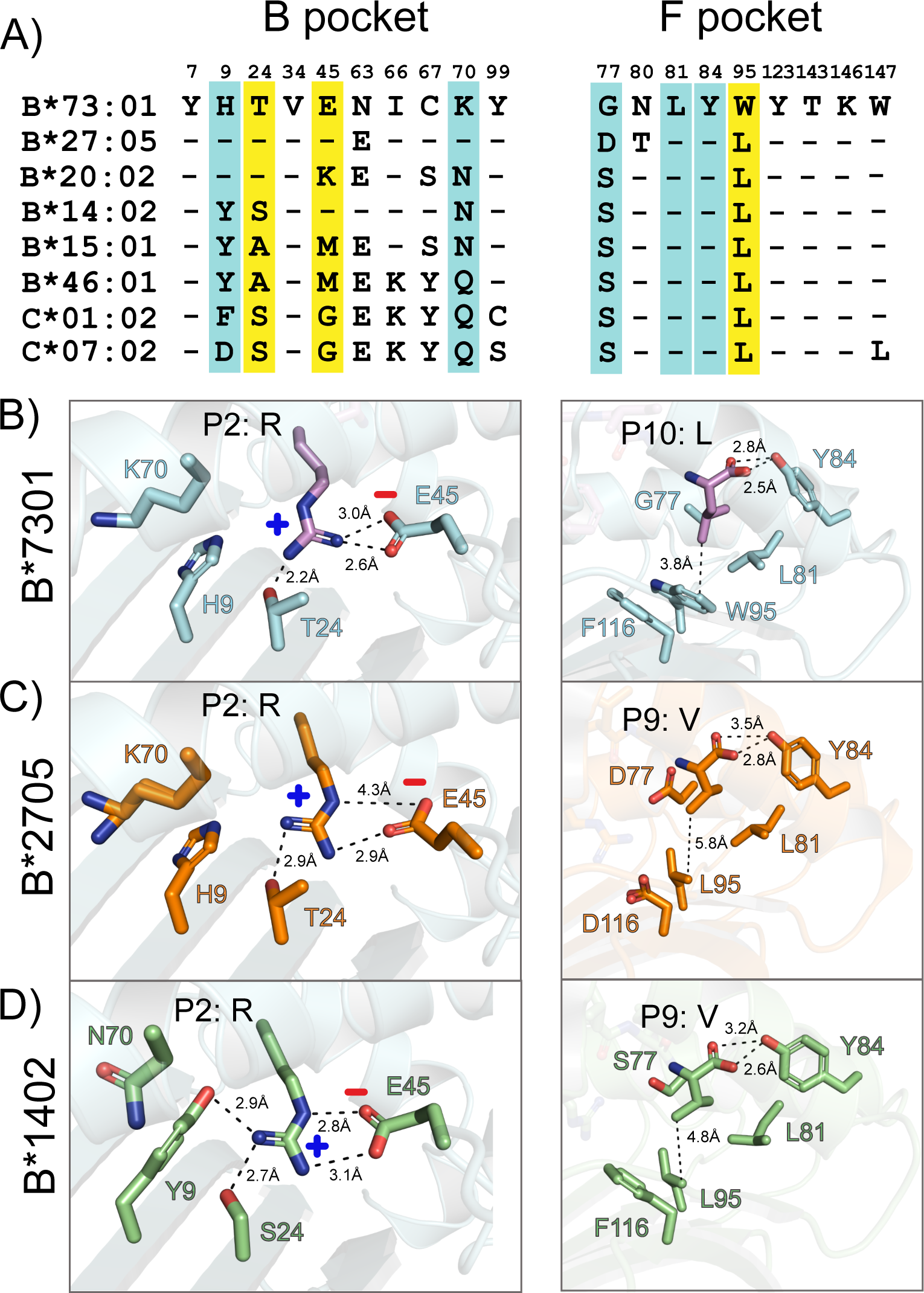
Variation in the B and F pockets of B*7301 provide flexibility in accommodating diverse peptide anchor residues. **A)** Residues lining the B (left) and F (right) pockets of B*73:01 and selected HLA alleles. Highlighted in yellow are key residues that engage the anchor residues shown in panels **B-D**, supporting residues are highlighted in cyan. Contacts between key peptide anchor residues and pocket residues in the crystal structures of B*73:01 **(B)**, B*27:01 **(C)** and B*14:02 **(D)**.

Although B*73:01 and B*27:05 have highly correlated peptide binding motifs, they present very few of the same peptide sequences when purified from the same cell. They share similar peptide anchor strategies within their B pockets, yet they have evolved quite different F pockets. One F pocket residue that is quite rare and carried by B*73:01 is W95, which only appears in roughly 12% of alleles in the set of alleles included in Sarkizova et al., while roughly 43% contain a leucine at this position (**Figure 4A**, right panel, highlighted in yellow). W95 likely contributes to the ability of B*73:01 to present peptides that have more diversity at the C-terminus than those presented by other HLAs likely due to its enhanced ability to shield small hydrophobic residues from water molecules, but also through CH/π interactions with proline residues at the C-terminus of the peptide (another preferred residue for B*73:01) (**Figure 4B**, right panel). This combination of relative constraint within the B pocket combined with hydrophobic plasticity within the F pocket represents a unique tradeoff for B*73:01 relative other alleles and contributes to its unique peptide repertoire, distinct from even B*27:05.

### The complex structure of B*73:01 and KIR2DL2 indicates the use of a unique docking angle

B*73:01 and B*46:01 are distinct from all other human B alleles in that they carry the C1 epitope (both V76 and N80 in the α1 helix) which confers reactivity to KIR2DL2/3. However, both alleles evolved their epitope independently (6, 15). Our complex crystal structure of B*73:01/KP1 with KIR2DL2 provides provides new insight into how these B alleles engage with receptors that are predominantly restricted to HLA-C alleles, specifically compared to the complex structures of 2DL2 with HLA-C*03:04 and HLA-C*07:02. Our complex shows the general binding mode of KIR2DL2 onto B*73:01 is grossly similar to that seen for other KIR complex structures published to date (**Figure 5A**). That is, the KIR binds directly over the F pocket of the heavy chain, making no contact with b_2_m with the D1 domain generally making contacts with the α 1 helix of B*73:01 and the D2 with the α 2 helix of the antigen binding region. Interestingly, however, when compared to other HLA/KIR2D complex structures (including KIR2DL2/ C*07:02, KIR2DL3/C*03:04, and KIR2DS2/A*11:01 (45, 46)) the docking angle of KIR2DL2 onto B*73:01 is shifted ∼15° towards the α 2 helix (**Figure 5B**). A contact map (**Figure 5C**) and buried surface area analysis (**Figure 5D**) of the intermolecular contacts between KIR2DL2 and B*73:01 compared with those of the complexes of HLA-C*07:02 and C*03:04 with KIR2DL2, show that the D1 domain of KIR2DL2 uses considerably fewer contacts (6 VDW and 1 h-bond) to engage the α 1 helix of B*73:01 than with the α 1 helices of HLA-C*07:02 (14 VDW, 3 h-bond, and 1 salt bridge) and C*03:04 (11 VDW, 2 h-bond, and 2 salt bridge). In contrast, the contacts between the D2 domain of KIR2DL2 and the α 2 helix of B*73:01 have slightly more VDW contacts, and share the same number of h-bonds (2) and salt bridges (3) as with the complexes with C*07:02 and C*03:04. Together, this results in an overall smaller footprint surface area for B*73:01 skewed towards the a2 helix, likely contributing to the “tilt” of KIR2DL2 on B*73:01 relative to these other complexes.

**Figure 5:**
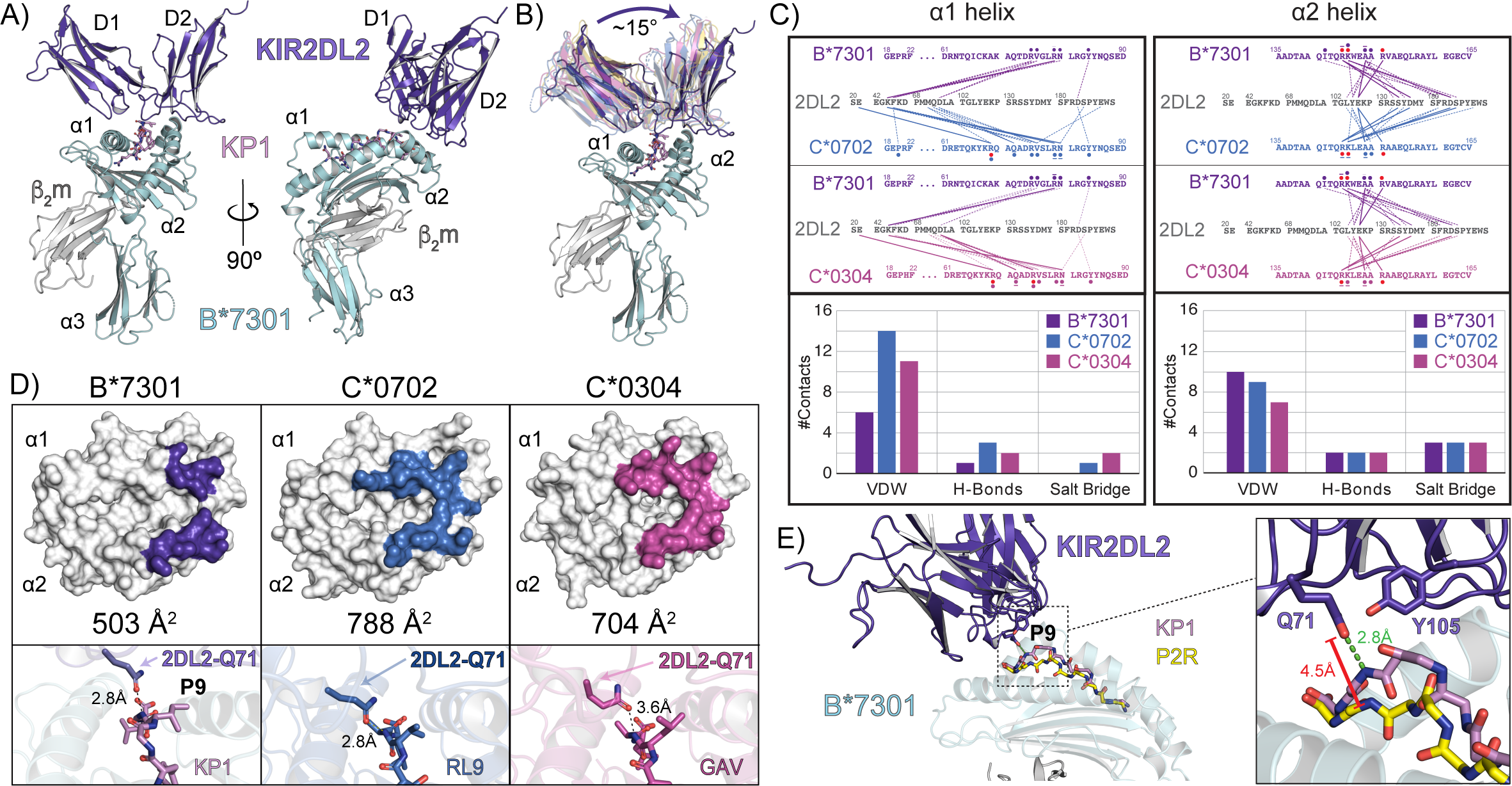
The three dimensional structure of the complex between B*73:01/KP1 and KIR2DL2. **A)** Two views, related by a 90 degree turn, of the complex; KIR2DL2 is shown in ribbon format, colored dark purple; HLA-B*73:01 is colored as in Figure 3: heavy chain in cyan, β_2_m in grey and KP1 peptide in mauve. KIR2DL2 engages B*73:01 directly over the F pocket. **B)** Comparison of the KIR2DL2/B*73:01-KP1 complex with that of representative KIR complexes with their cognate HLA. Shown are KIR2DL2 and KIR2DL3 engagement of HLA C*07:02 (PDB ID: 6PA1 and 6PA2; semi-transparent marine blue and semi-transparent magenta, respectively) and KIR2DS2 engagement of HLA A*11:01 (PDB ID: 4N8V, semi-transparent yellow) (16, 45). HLAs were superimposed based on their Cα backbones. Shown is the angle differences between the KIR2DL2 docking on B*73:01 in comparison with the KIRs docking on C*07:02 and A*11:01. **C)** Contact analysis between the α helices of B*73:01, C*07:02 and C*03:04 and KIR2DL2. Shown are the amino acid sequences of the α1 helix (left) and α2 helix (right) of the HLAs. Lines denote contacts between KIR2DL2 and these HLAs; solid lines are hydrogen bonds (<3.4 Å) and dashed lines are van der Waals (<4.5 Å) coloring is based on the relevant HLA. Symbols above and below the amino acid sequences indicate which type of interaction each residue contributes to: circles in backbone color: VDW, lines: h-bonds, red circles: electrostatic/salt bridges. Below the contact figure are bar graphs presenting the total number of contacts of each type per complex. **D)** The KIR2DL2 binding footprint on B*73:01 (dark purple) versus C*07:02 (marine blue, PDB ID: 6PA1) and C*03:04 (magenta, PDB ID: 1EFX). HLAs are shown in surface representation colored white with the relevant KIR2DL2 buried surface area footprints colored as indicated above. Below each footprint representation is a zoomed view of the key KIR2DL2 residue Q71 in the interface with the respective HLAs. Q71 forms a hydrogen bond with the P8 position of the presented peptide backbone. In the complex structure with B*73:01 (left panel), Q71 adopts a different conformation than it does when engaging C*07:02 (middle) and C* 03:04 (right). **E)** Comparison between the complex between KIR2DL2 and B*73:01 with the KP1 peptide (10mer) and the modeled complex with the P2R peptide (9mer). Distance measurements are between the Q71 sidechain of 2DL2 and the P9 mainchain nitrogen of each peptide.

A close-up view of the buried surface area between the KIR and presented peptide backbones shows that Q71 of KIR2DL2 has a direct h-bond contact with the main chain nitrogen of P9 of the KP1 peptide, either a result, or cause of, the skewed KIR binding (**Figure 5D**). In other complex structures, Q71 remains closer to the α 1 helices of their corresponding HLA ligands. Because the KP1 peptide is likely presented this way regardless of binding to KIR2DL2, the engagement of Q71 with the backbone of KP1 likely further contributes to the unique docking angle of KIR2DL2 onto B*73:01. Modeling of KIR 2DL2 onto the nonamer P2R peptide/B*73:01 complex structure provides some insight into how peptide length and sequence may regulate KIR2DL2 engagement in a peptide specific way. Whereas KIR2DL2 via Q71 forms a strong h-bond with the KP1 backbone at position 9 (distance ∼ 2.8Å), the backbone of P2R is ∼4.5Å away from Q71 (**Figure 5E**), too far a distance for effective h-bonding. KIR2DL2 also has an additional VDWs contact with KP1 via Y105, which is not resolved in our P2R peptide complex model. This complex structure also suggests that for KIR2DL2 engagement, peptides presented by B73 need to be within a narrow length range; too short, and Q71 cannot engage, whereas if they are too long (i.e. greater than 11mers), then they may disrupt KIR2DL2 binding by sterically inhibiting its binding over the F-pocket, as these peptides appear to be accommodated through “bulging out” near the C-terminus.

## Discussion

Our results demonstrate that B*73:01, an archaic allele that was likely re-introduced to the human population through introgression, presents a restricted repertoire of unusually long peptides with a preference for arginine at P2 and a hydrophobic residue at the C-terminal position. While other HLA alleles exist with these preferences (i.e. B*27:05, B*07:02 and B*14:02, **Figure 1**), the B*73:01 peptide repertoire is highly unique, likely the result of a combination of factors, one of which is the ability to present longer peptides 11+ amino acids in length. Our structural analysis of B*73:01 with the KP1 10-mer peptide provides a model for understanding the molecular basis for this repertoire selection including a shift in the N-terminal peptide anchoring (**Figure 3C, D**) and a skewing of the peptide “bulge” towards the C-terminus. While all these factors combine to generate a binding cavity that is unlike that of many other HLAs, there are also other factors, such as availability of peptides or distinct interactions of B*73:01 with the peptide-loading complex (PLC) machinery that may also play a role in restricting this repertoire. In comparing the diversity of nonamers at each position more broadly, we noted that B*27:05 has even greater restriction at P2 than B*73:01 (**Figure S1D**). It is known that the KK10 peptide bound by B*27:05 of HIV origin shows signatures of escape mutations at the P2 position, mutating an arginine anchor to a lysine in order to escape B*27:05-mediated T-cell surveillance (47). More diversity in the B*73:01 P2 anchor position may allow B*73:01 to be more tolerant of such escape mutations relative to B*27:05. Indeed, our data suggest that 1% of B*73:01 presented peptides contain a lysine at P2, whereas only 0.2% of such peptides bind to B*27:05.

Highly compact peptidomes, like that of B*73:01, are thought to correlate with protection against closely related pathogens (36). To our knowledge, there are no epidemiologic studies that have correlated B*73:01 with the development or outcome of infectious disease, likely due to the low frequency of B*73:01 in the modern human population. However, HLA alleles that encode the C1 epitope, in combination with KIR2DL3, are known to provide protection against hepatitis C virus (18) (an RNA virus), providing some rational for B*73:01 having this KIR specificity. Our data shows that B*73:01 is expressed at high levels on the cell surface, presumably much higher than HLA-C alleles, so its carrying of the C1 epitope may provide a strong signal to inhibitory KIR2DL2 and KIR2DL3. Our structural data provides insight into how an HLA-C biased KIR can engage with a C1 epitope carrying HLA-B allele; B*73:01 engages KIR2DL2 with a skewed footprint, in part due to the engagement of the peptide backbone mediated by Q71 of 2DL2 (**Figure 5C**). Because peptides presented by B*73:01 appear to selectively “bulge” out from the binding groove near the C terminus of the peptide, the longer the peptide the more likely it will sterically hinder the engagement of these KIR2D receptors. Presentation of longer peptides from pathogens might thus both recruit a T cell response (not investigated in this report) and disrupt any NK cell inhibition mediated through the C1 epitope of B*73:01, leading to a more robust pathogen-specific response. Intriguingly, and in support of our hypothesis, a recent study suggests that RNA viruses were likely the major drivers of adaptive introgression of archaic alleles from Neanderthals to modern Europeans (48). Of course, as is the case with B*27:05, it is also likely that predisposition to autoimmune disease may also result. This is one possible explanation for the paradoxical low incidence of B*73:01 in the modern human population: as a result of its unique characteristics, it may have conferred exceptional protection against disease, but also exceptional predisposition to autoimmunity (presumably one under selective pressure) and thus fallen outside of the normal range for utility of an HLA molecule. Intriguingly, there is one case report in which B73 is strongly linked to pediatric mycosis fungicides (49) although the authors rightly conclude that the numbers are too low for the finding to be conclusive in any way.

In summary, our study shows that archaic B*73:01 is a highly expressed HLA with a unique peptidome of unusually long length which may modulate its ability to engage with the KIR2Ds through its C1 epitope. These structural and functional characteristics distinguish B*73:01 from other HLA class I alleles and likely provided early modern human migrants that inherited this allele from archaic humans with a selective advantage as they colonized Europe and Asia.

## Experimental Procedures

### Peptidome sequencing and analysis

Isolation and analysis of peptides was performed as previously described (15). Briefly, 721.221 cells were transfected with constructs encoding a soluble form of HLA-B*73:01. The sHLA-peptide complexes were affinity purified from the cell-supernatant on an anti-W6/32 Sepharose column, eluted and the bound peptides dissociated from the HLA by denaturation. Peptides were separated from the denatured proteins by ultrafiltration and separated into ∼40 fractions by reverse-phase high-performance liquid chromatography (HPLC) as described. Approximately 25% of each HPLC fraction was injected into a nano-scale reverse phase liquid chromatography Eksigent nano-LC-4000 (Sciex) system. Column specifications, mobile phase solvents, and the elution gradient were as described (50). Eluted fractions were ionized using a NanoSpray III (Sciex) ion source into a Sciex TripleTOF 5600 mass spectrometer collecting LCMS spectra in DDA mode. Sequences were assigned to spectra using PEAKS 7 at a 1% FDR as described in (50).

For each amino acid in a peptide, four biochemical properties (molecular weight, hydropathy index, surface area and isoelectric point) were determined. Thus 36 variables were generated for each nonamer peptide. Peptides of length 8-15 were extracted from the eluted ligand data for the four HLA class I alleles. A GibbsCluster analysis was performed on each data set to identify the majority binding motif and remove noise in the data.

### Integration with and analysis of Sarkizova et al. data

Curated peptide lists were downloaded from ftp://massive.ucsd.edu/MSV000084172/ and compared directly to peptides eluted from B*73:01. To calculate correlation coefficients in peptide space between all alleles, amino acid occurrences of all 20 amino acids were calculated for each position of all nonamers generating a 9x20 matrix for each allele. These matrices were then flattened into vectors of 180 integers for each allele and used to directly calculate the Pearson correlation coefficients between alleles of presented nonamers. For plotting, alleles were clustered by hierarchical clustering implemented by the ggcorrplot function of the ggcorrplot package in R.

To calculate correlation coefficients between alleles in pocket space, pseudosequences for each allele were used as described in (51). In brief, following a multiple sequence alignment, positions within the alignment that were 100% conserved between alleles were removed, leaving only variable aligned columns. These sequences were then used to calculate 61 different descriptors of their amino acid sequences using the aaDescriptors function of the Peptides package (https://pypi.org/project/peptides/) and default parameters. This generated a vector of 61 values for each allele which was then used to calculate Pearson correlation coefficients between all alleles. For plotting, alleles were clustered by hierarchical clustering implemented by the ggcorrplot function of the ggcorrplot package in R (https://cran.r-project.org/web/packages/ggcorrplot/readme/README.html).

For plotting sequence logos, the R package ggseqlogo was used (52, 53). For calculating Shannon and Simpson diversity indices, the diversity function from the R package vegan (53) was used.

### Cell-surface expression experiments

721.221 cells, which lack endogenous HLA class I expression, were transfected with constructs encoding B*73:01, B*46:01, B*15:01 or C*01:02 and cultured as previously described. Cells expressing HLA class I were detected by flow cytometry (Accuri C6 cytometer, BD Biosciences). Expression levels of each allele was determined from the median fluorescence intensity (MFI) of the W6/32 antibody bound to each positive staining cell. Three independent transfections with at least 50,000 cells each were performed for each allele tested.

We examined the cell-surface expression of wild-type and KYR/ICA mutant 3x FLAG-tagged B*73:01 and B*46:01 in HeLa cells. Recombinant cDNA encoding amino acids 1-338 of B*73:01 and B*46:01 with an N-terminal 3X FLAG-tag were manufactured by Genscript (Piscataway, NJ). Site-directed mutagenesis was performed with the QuikChange Kit (Stratagene), according to the manufacturer’s instructions, to generate the two swap KYR/ICA mutants. HeLa cells were transfected with these constructs using the Fugene transfection reagent (Promega) and cultured as previously described (54). Cells expressing FLAG-tagged HLA class I were detected by flow cytometry (Accuri C6 cytometer, BD Biosciences). Expression levels of each allele or mutant were determined from the MFI of FITC-conjugated anti-FLAG antibody bound to each positive staining cell. Three independent transfections with at least 50,000 cells each were performed for each allele tested.

### Phage display selection protocol

To obtain high-affinity binders, five rounds of selection were conducted using the phage display selection protocol previously described (55). Biotinylated HLA-B*73:01 was immobilized onto streptavidin-coated paramagnetic beads (Promega) for the selection process. In the initial round, 500 nM of HLA-B*73:01 was immobilized on 200 μL SA magnetic beads and incubated with 1 mL phage library (10^10 CFU) for 1 hour at room temperature with gentle agitation. The beads were subjected to three washing cycles to remove non-specific phage, then introduced to log-phase E. coli XL-1 blue cells and incubated for 20 minutes at room temperature. Subsequently, media containing 100 μg/mL ampicillin and 10^9 p.f.u./mL of M13K07 helper phage (NEB) was added for overnight phage amplification at 37°C. The amplified phage was precipitated in 20% PEG/2.5 M NaCl for 20 minutes on ice in preparation for subsequent rounds. Prior to each round, the phage pool underwent negative selection against empty paramagnetic beads for 30 minutes with shaking to eliminate non-specific binders. The antigen concentration was systematically reduced from 500 nM to 10 nM from the first to the fifth round (2nd round: 250 nM, 3rd round: 100 nM, 4th round: 50 nM, and 5th round: 10 nM). After phage binding, the beads were subjected to five washing rounds with 0.5% BSA/PBST. Bound phages were eluted using 0.1 M glycine, pH 2.6, and neutralized with TRIS-HCl, pH 8. The phage eluate was then used for E. coli infection and phage amplification as previously described. Following the fourth and fifth rounds, infected cells were plated on ampicillin agar, and 192 colonies were selected to produce phage clones for single-point phage ELISA assay. Promising clones demonstrating high specificity were sequenced, reformatted into an RH2.2 expressing vector, and produced as previously described (56).

### Phage Enzyme-Linked Immunosorbent Assays (ELISA)

HLA-B*73:01, at a concentration of 50 nM, was directly immobilized onto high-binding experimental wells (Greiner Bio) for a duration of 30 minutes. Subsequently, the wells were subjected to extensive blocking with 2% BSA for 1 hour to minimize non-specific binding. Following a 15-minute incubation period with phage, the wells underwent a rigorous washing protocol, consisting of three cycles with 0.5% BSA/PBST. The wells were then incubated with Protein L-HRP (Thermo Scientific) at a 1:5000 dilution in HBST for 20 minutes. After another thorough washing step, the plates were developed using TMB substrate (Thermo Scientific). The enzymatic reaction was quenched with 10% H3PO4, and the absorbance was quantified spectrophotometrically at 450 nm (A450).

## Structural determination and analysis by electron microscopy

### Sample preparation and data acquisition

Quantifoil (R1.2/1.3, 200 mesh) gold grids (Ted Pella) were glow-discharged for 10 seconds at 20 watts using the Solarus 950 Plasma Cleaner System (Gatan). Fluorinated octyl maltoside (Anatrace) was added to the B*7301-B.1-B.8 complex to a final concentration of about 0.07 mM before plunge freezing. Sample (3 ul at 0.6 mg ml-1) was applied to grids, which were blotted for one second at blot force two, using a Vitrobot Mark IV (ThermoFisher) maintained at 8°C in a 100% humidity environment, and plunge frozen into liquid ethane. Data were collected using a 300 kV Titan Krios G3i (ThermoFisher) microscope equipped with a K3 detector (Gatan). We used a nominal magnification of 81,000x, translating to a pixel size 0.5325 Å in the raw micrographs. EPU software was set to automated acquisition mode and collected 5291 movies, each with 50 frames, subject to a total dose of 60 e-/Å^2^.

### Image processing

Movies were motion-corrected and dose-weighted using MotionCorr2 (57) in the RELION-3.1.3 (58) wrapper, with data binned 2X to 1.065 Å/pixel. Micrographs were imported into cryoSPARC v3.3.1 (59) and subjected to CTF Patch Estimation. Particle coordinates were identified by reference-free “blob” particle picking. A 256 pixel box was used to extract 4X binned (2.13 Å/pixel) particles, which were then sorted into fifty 2D classes, several of which were selected as templates. Template particle picking yielded an initial set of 4,007,333 particles, which were culled to 1,531,860 following an additional round of 2D classification and used to generate three ab initio 3D volumes. These volumes were used as templates to sort particles into three Heterogeneous Refinement classes. Particles from the two best classes were subjected to another round of 2D Classification, from which four 2D classes were chosen for a new round of template particle picking.

4,844,663 particles were extracted and sorted into 50 classes in two consecutive rounds of 2D Classification. 2,217,300 particles were extracted in 256 pixel boxes (at 2.13 Å/pixel) and selected for Ab-Initio Reconstruction into four classes. The two highest-resolution 3D volumes, as well as one junk (“garbage collector”) class were selected as initial models for Heterogeneous Refinement. All three output volumes, as well as 1,141,440 particles associated with the highest-resolution class, were selected for an additional round of Heterogeneous Refinement. The highest-resolution refinement map and 732,360 associated particles were then selected as inputs for Homogeneous Refinement. Though estimated to have 3.2 Å resolution by cryoSPARC, the Homogeneous Refinement map exhibited prominent streaking in underrepresented orientations. The map and particles were therefore exported to RELION-3.1.3 for further processing.

Following 2D Classification of the exported particles, 682,726 were subjected to a round of high-iteration 3D Classification in which they were aligned to the 3D volume obtained from cryoSPARC’s Homogeneous Refinement and sorted into two classes. The angular sampling interval was progressively lowered every 50 iterations. The final set of 502,247 particles underwent several rounds of 3D auto-refinement, as well as CTF refinement. A solvent mask was generated for the highest-resolution auto-refined map, which then underwent B-factor sharpening to yield the final reconstruction.

### Refinement

Initial models used included the crystal structure of B*7301 determined in this study (PDB: 8TMU) and AlphaFold2 (60) models generated for B.1 and B.8 Fabs using their respective primary amino acid sequences. Initial model docking into the highest-resolution refined map was done manually using ChimeraX (61) version 1.5 and the *fitmap* command. Additional rounds of real-space refinement were performed using PHENIX (62) version 1.20.1-4487-000.

### Crystal structure determination and analysis

Soluble B*73:01 heavy chain (residues 1-276) and β_2_M light chain constructs were expressed in E. coli and refolded from inclusion body preps with respective peptides. A soluble KIR2DL2*001 construct (residues 1-225 of the mature protein), covalently linked to an N-terminal 8x HisTag, separated by a 3C protease cleavage site, was expressed in BTI-Tn-5B1-4 insect cells (High Five) cells cultured in Insect-XPRESS Protein-free Insect Cell Medium (Lonza). BestBac 2.0 linearized DNA (Expression Systems) was used to transfect Sf9 cells to generate a P1 viral stock. Virus was then amplified by shaking culture in Sf9 cells. Following Ni-NTA affinity purification, KIR2DL2 was treated for 2 hours at 37°C with EndoF (made recombinantly in the Adams Lab) prior to another round of Ni-NTA affinity purification to remove any non-HisTagged EndoF. The 8xHisTag was removed using 3C protease (made recombinantly in the Adams Lab). Recombinant B*73:01 and KIR2DL2 was purified separately by SEC into 10 mM HEPES, 150 mM NaCl, pH 7.2 (HBS), mixed at a 1:1 molar ratio, and concentrated to 6-10 mg/mL. Crystals were initially grown in less than 24 hours in mother liquor containing Tris-HCl pH 8.5 and 25% PEG 8000. These crystals were then crushed and used as microseeds to grow a larger crystal in mother liquor containing Tris-HCl pH 8.75 and 20% PEG 8000, which took 1-2 weeks to grow.

CCP4, Coot, and PHENIX software suites were used for molecular replacement and refinement. PyMol was used to make figures and measure distances between atoms (58, 59, 63-67).

## Data availability

The refined structure coordinates have been deposited in the Protein Data Bank (www.rcsb.org) with accession code 8TNJ for the cryo-EM structure and 8TMU for the crystal structure.

## Supporting information

B73-peptidome

## Acknowledgements

This work was completed with resources provided by the University of Chicago’s Research Computing Center and resources of the Advanced Photon Source, a U.S. Department of Energy (DOE) Office of Science user facility operated for the DOE Office of Science by Argonne National Laboratory under Contract No. DE-AC02-06CH11357. We would like to thank the University of Chicago Cytometry and Antibody Technology Core Facility (RRID: SCR_017760) and Dr. James Fuller of the Advanced Electron Microscopy Core Facility (RRID:SCR_019198) for maintaining critical instruments and providing technical expertise during experiment planning, data collection and analysis.

## CRediT author statement

**Philipp Ross:** Writing - Original Draft, Visualization, Investigation, Formal analysis. **Hugo G. Hilton:** Conceptualization, Writing - Original Draft, Investigation, Formal analysis. **Jane Lodwick:** Investigation, Formal analysis. **Tomasz Slezak:** Investigation, Formal analysis. **Lisbeth A. Guethlein:** Investigation, Writing - Review & Editing. **Curtis P. McMurtrey:** Investigation, Formal analysis. **Alex S. Han:** Investigation. **Morten Nielsen:** Writing - Review & Editing. **Daniel Yong:** Investigation. **Charles L. Dulberger:** Investigation. **Kristof T. Nolan:** Investigation, Formal analysis. **Sobhan Roy:** Investigation, Formal analysis. **Caitlin D. Castro:** Writing - Review & Editing. **William H. Hildebrand:** Funding acquisition, Writing - Review & Editing**. Minglei Zhao:** Funding acquisition. **Anthony Kossiakoff:** Funding acquisition. **Peter Parham:** Conceptualization, Supervision, Funding acquisition. **Erin J. Adams:** Writing - Original Draft, Visualization, Supervision, Funding acquisition

## Funding and additional information

This work was supported by the NIH under the following grant numbers: R35GM143052 to M.Z., GM117372 to A.K., AI22039 and AI17892 to P.P., and AI155984 to E.J.A.

## Supplementary Materials

**Figure S1:**
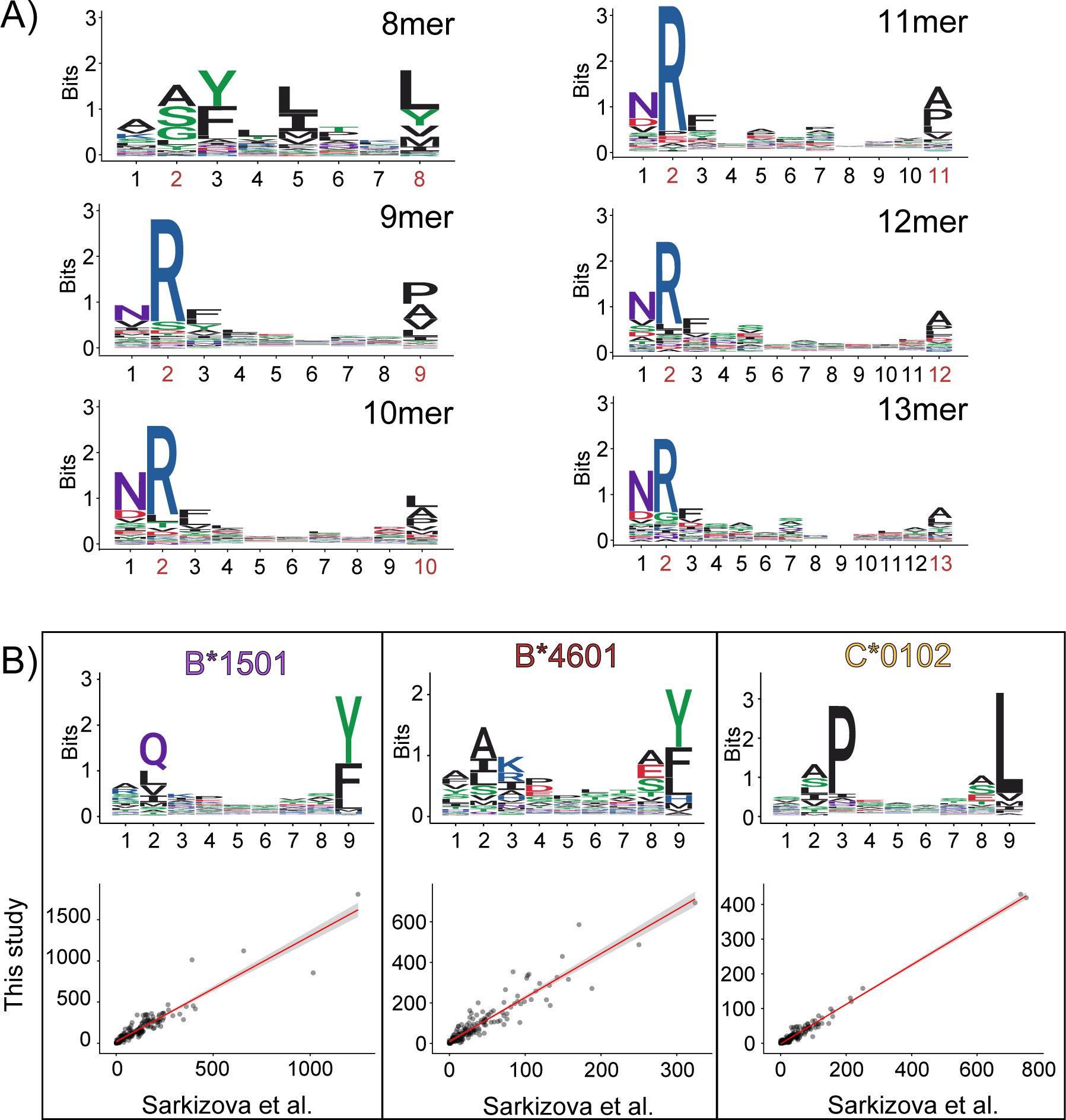
**A)** Sequence logos of peptides eluted from B*7301 stratified by length including peptides of length 8 (8mer) to length 13 (13mer). The colors of each amino acid correspond to their biochemical characteristics: acidic (red), basic (blue), hydrophobic (black) and polar (green). **B)** The 9mer peptide binding motif profiles (upper panels) of B*1501, B*4601, and C*0102 from Sarkizova et al. correlate positively (Pearson) with data generated in our previous study as shown through scatterplots (lower panels) of total bound peptides by alleles in this study plotted across the cell-surface expression values of the same alleles as expressed on 721.221 cells taken from Bashirova et al. 2020. Correlations were calculated using the *cor.test()* function in R and plotted using the *lm()* function.

**Figure S2:**
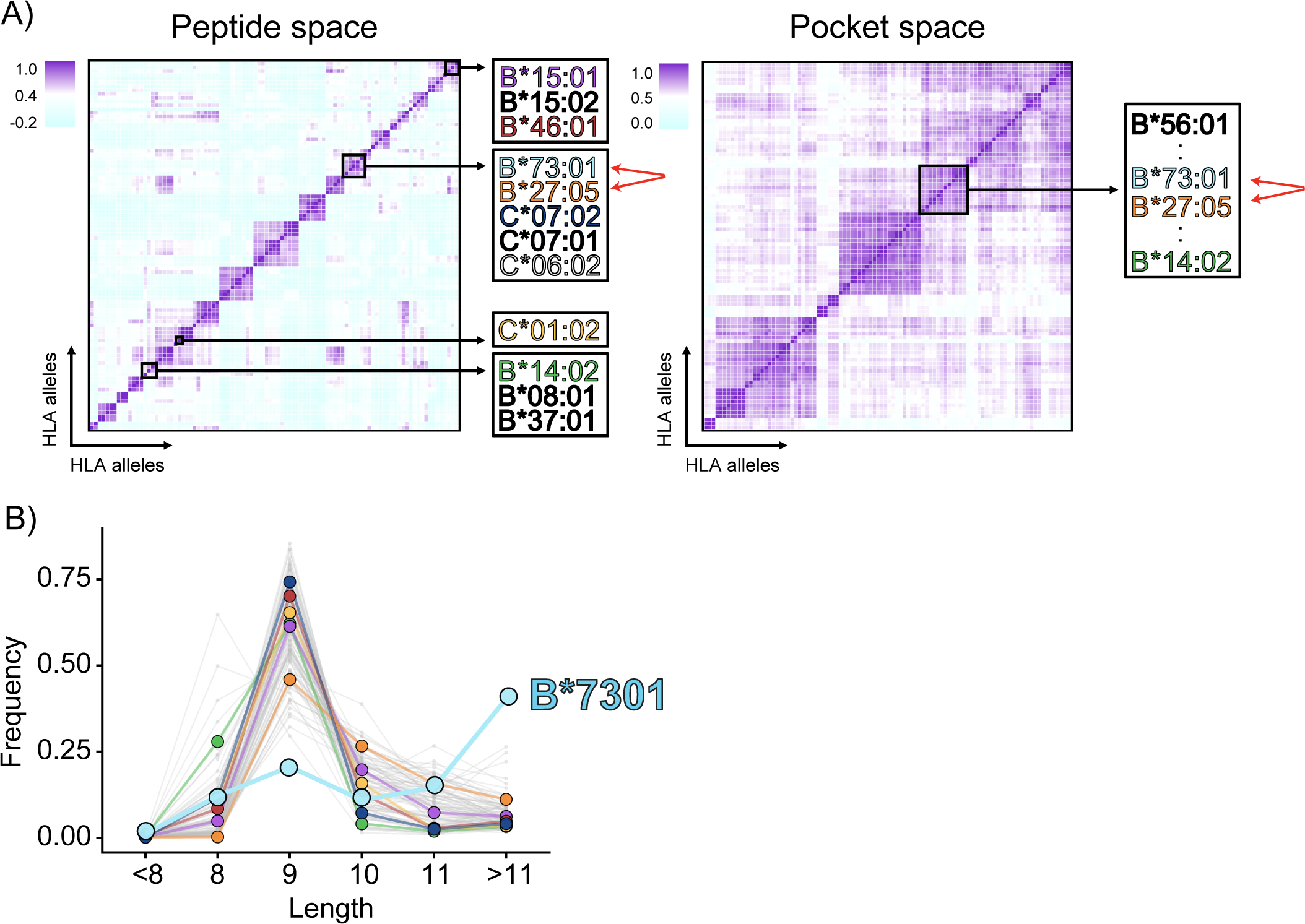
**A)** Heatmaps were generated using data from Sarkizova et al. was combined with eluted peptides from HLA-B*73:01 and representative peptides known to bind HLA-E (IEDB references 1037304, 1032571, and 1004539) and then correlated (Pearson) with other peptidomes (left; peptide space) and other MHC pocket residues (right; pocket space), then clustered based on similarity to highly alleles with similar peptide binding motifs and/or similar binding grooves. Alleles with similar peptide repertoires and/or similar binding grooves are depicted on the right. **B)** Length distributions of different alleles from Sarkizova et al. grouped into peptides less than 8, 8, 9, 10, 11, or more than 11 residues in length and including B*7301, shown in cyan.

**Figure S3:**
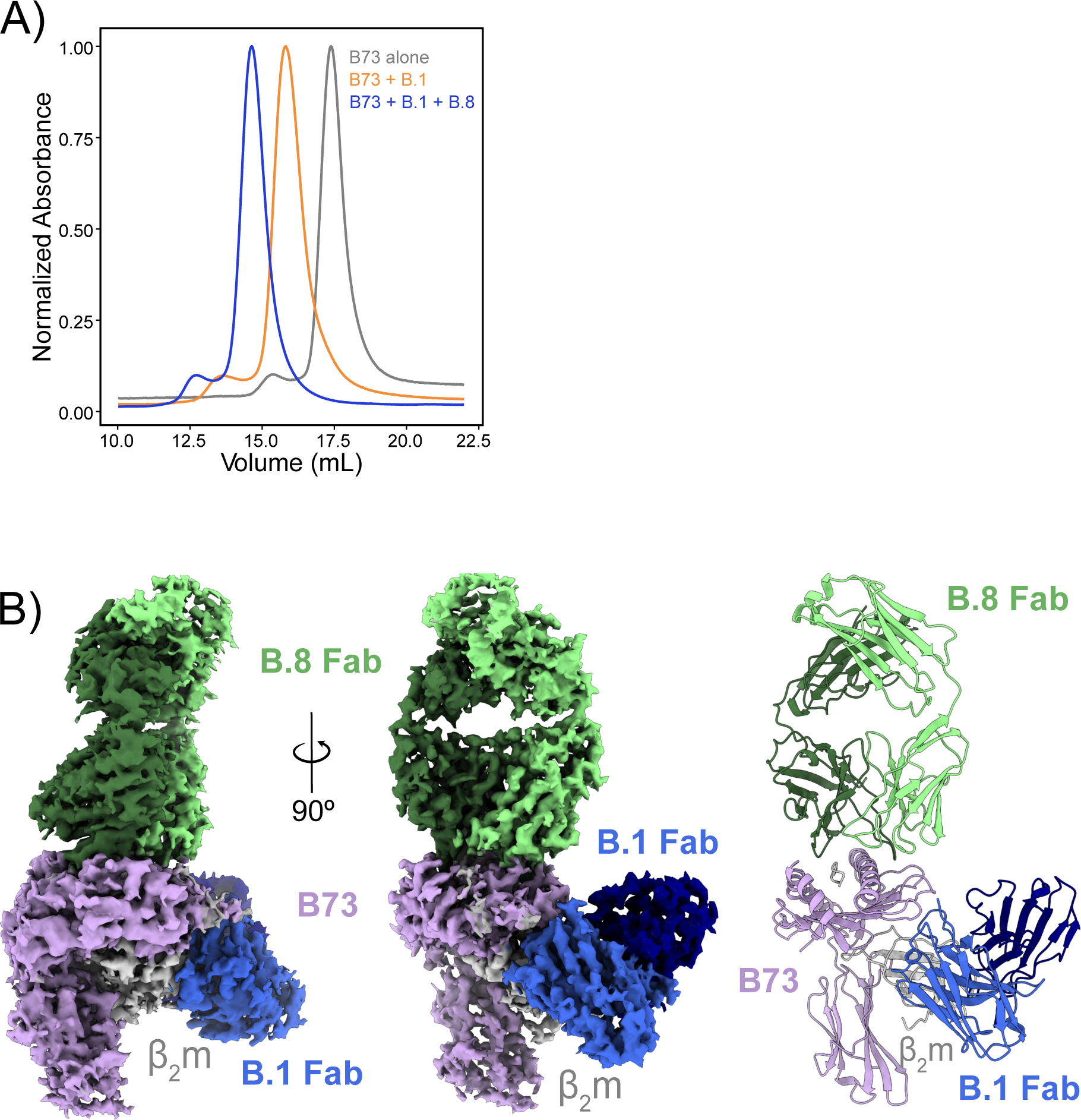
**A)** Size exclusion chromatography curves showing elution of HLA-B*73:01 following injection of the HLA by itself, with only the B.1 Fab, or with the B.8 and B.1 Fabs. **B)** The masked density map at a threshold of 0.02 of HLA-B*73:01 bound to the P2R peptide and complexed with the B.8 and B.1 Fabs.

**Figure S4:**
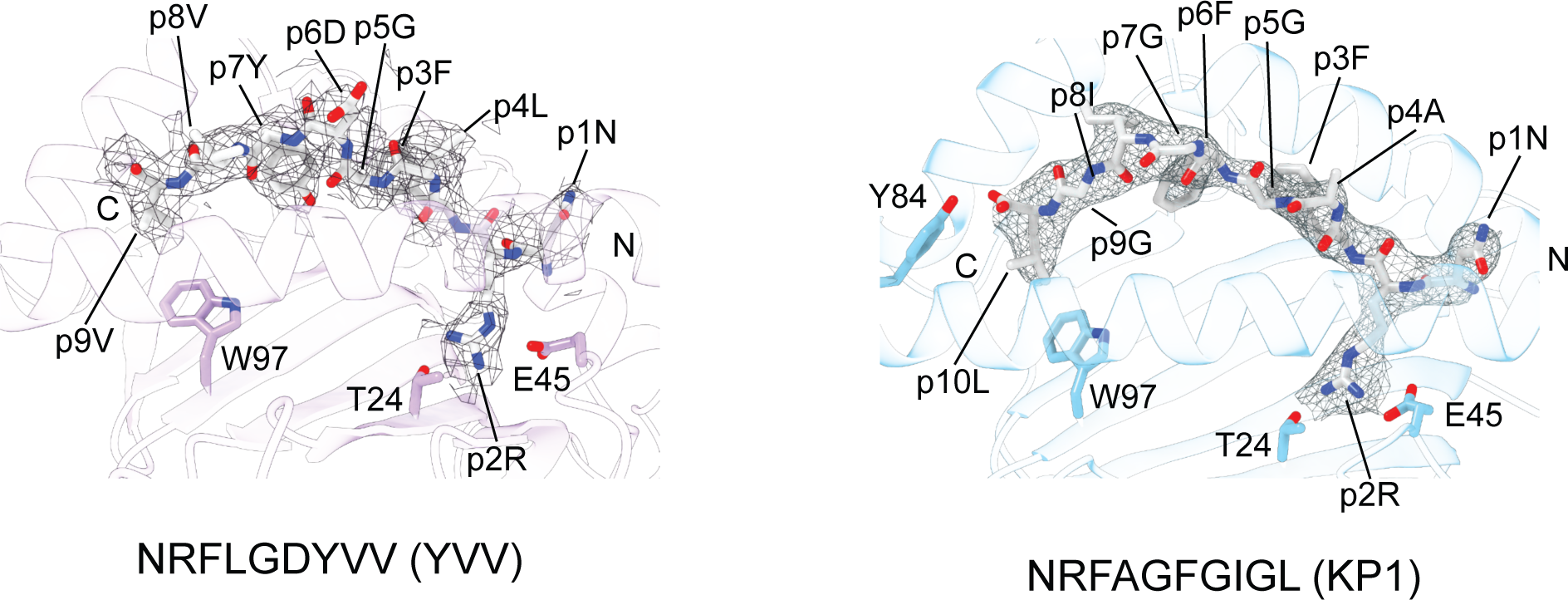
The mesh surrounding the P2R peptide for the CryoEM (left) structure, representative of the masked map at a threshold of 0.02. The mesh surrounding the KP1 peptide for the X-ray crystal (right) structure, representative of the 2Fo-Fc map restricted to a certain radius surrounding only the peptide itself.

**Figure S5:**
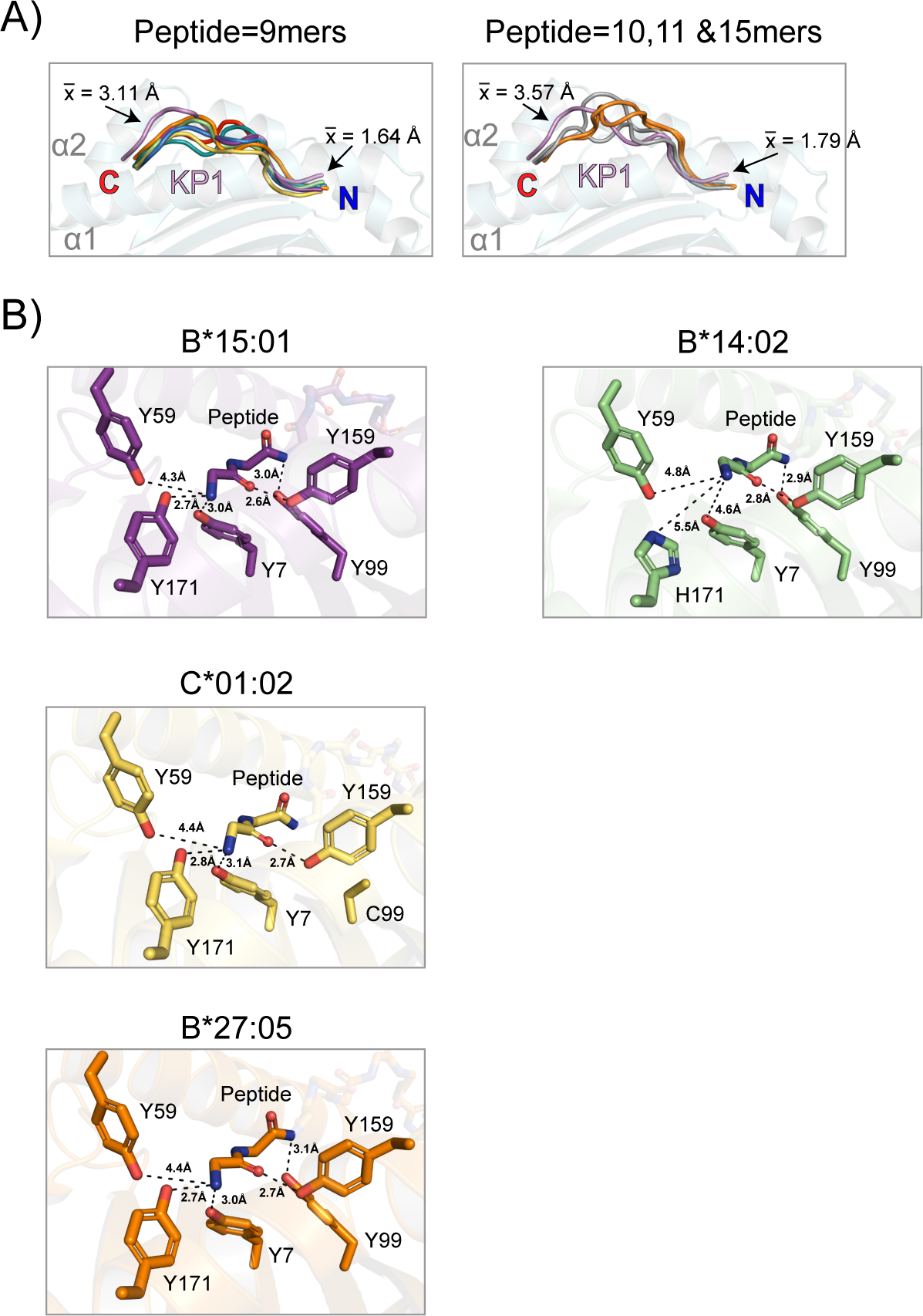
**A)** KP1 bulges out at the C-terminal end of the peptide over the F pocket of B*7301 in a manner not seen in other alleles presenting a variety of 9mers. Colors correspond to allele colors as in Figure 1 and Figure S1. Distances were calculated in PyMol as the distance between alpha carbons at the second to last or first residue within each peptide ligand. Comparing even longer peptides, KP1 still stands out. Distances were calculated as in panel D. **B)** A close up of A pockets are shown for alleles B*1501, B*2705, C*0102, and B*1402. Distances were calculated using PyMol.

**Figure S6:**
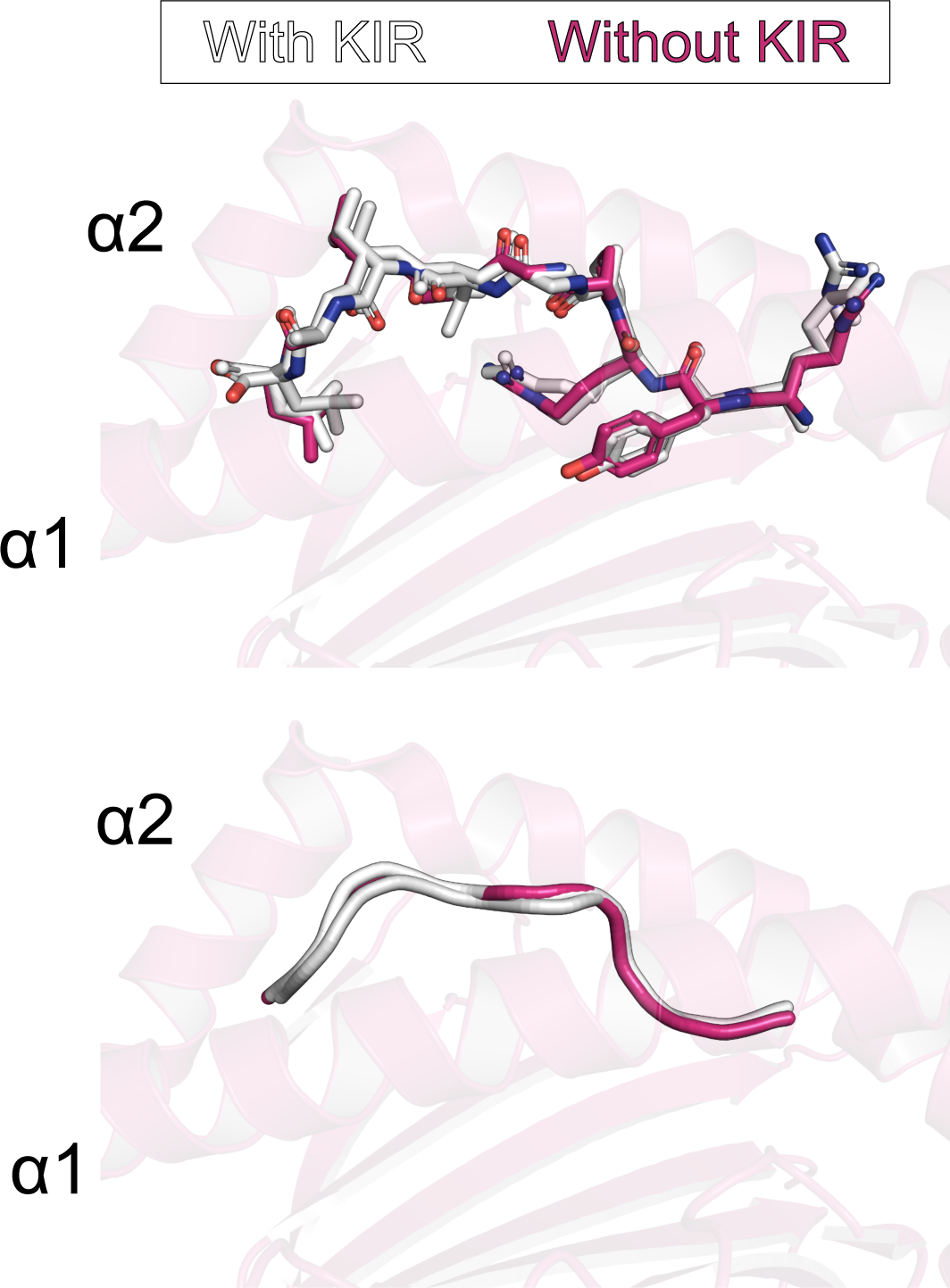
Three structures of C*0702 were aligned using β_2_m as a reference. In magenta is a structure of C*0702 presenting RYRPGTVAL without a bound KIR. Two more structures in white are structures of C*0702 presenting the same peptide, bound to KIR2DL2 or KIR2DL3.

**Figure S7:**
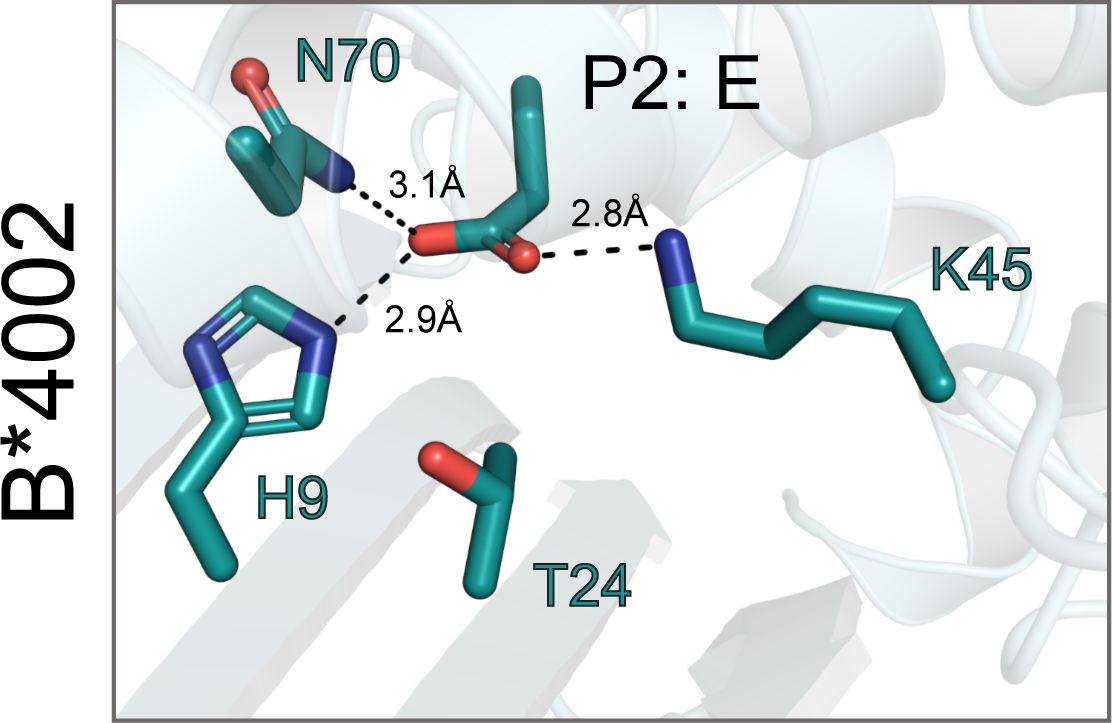
A close up of B pocket of B*4002 show how it uses similar and different residues located on different sides of the binding pocket to accommodate positively or negatively charged anchors.

**Table S1.**
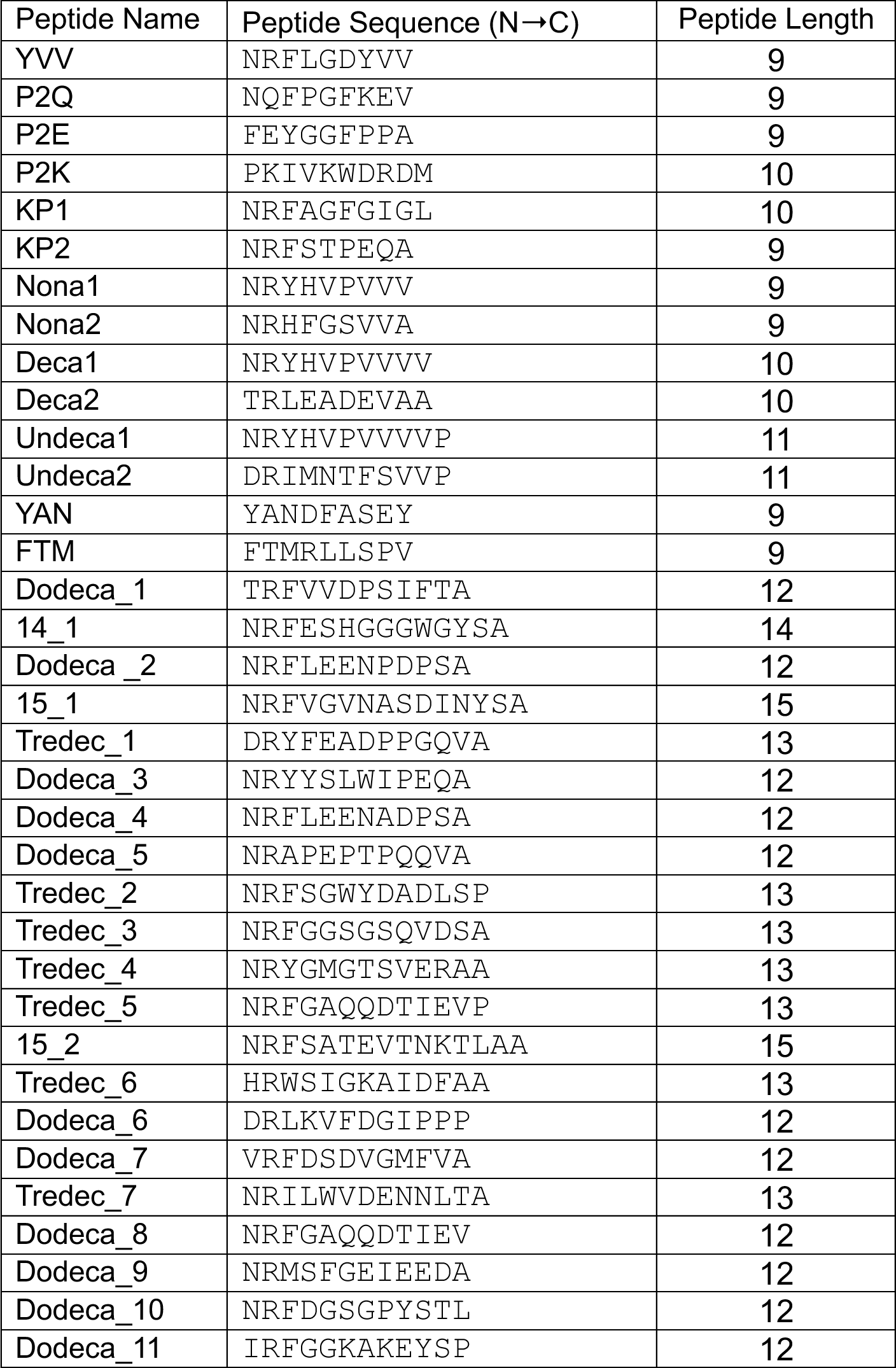
Peptides eludes from B73.

**Table S2:**
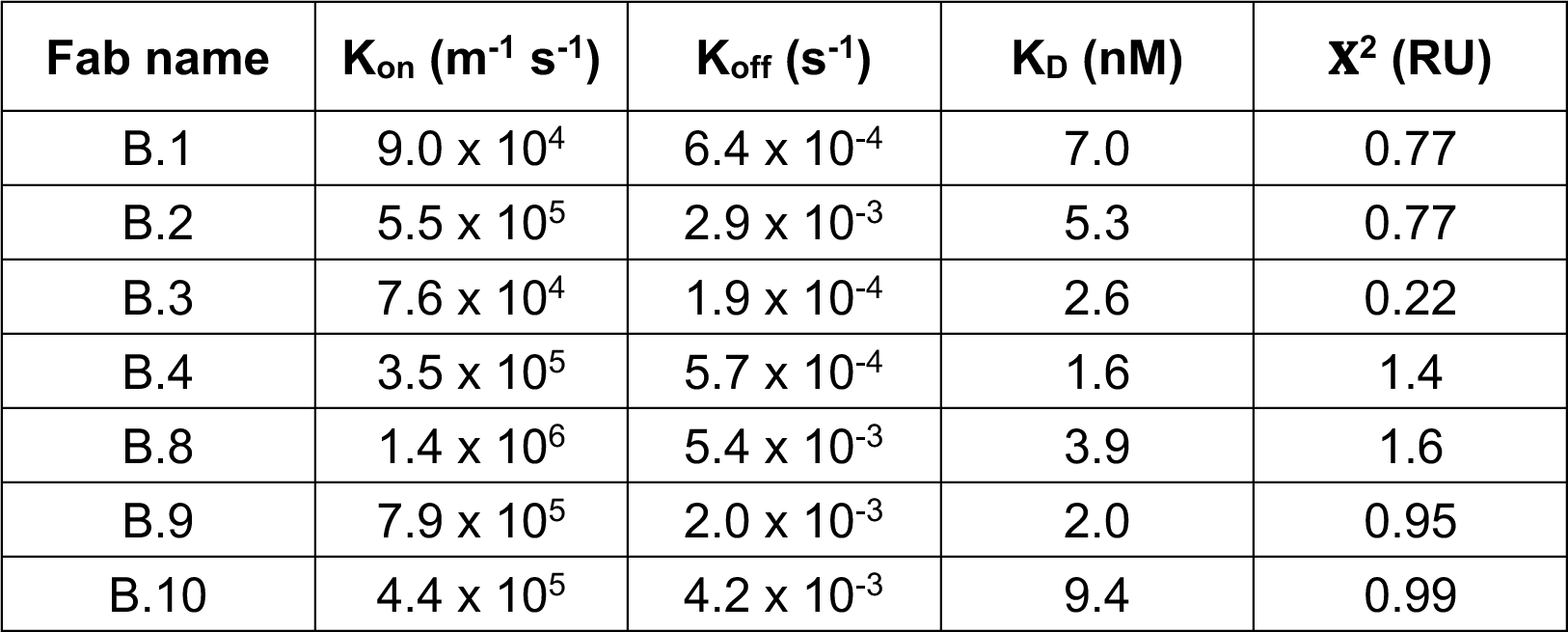
Kinetics of B73 specific Fabs.

**Table S3.** Data collection, processing and refinement statistics for B.1-B.8-YVV-B2M-B73.

|  |  |
| --- | --- |
| **Microscope** | Titan Krios |
| **Nominal magnification** | 81,000 |
| **Voltage (kV)** | 300 |
| **Electron exposure (e/Å<sup>2</sup>)** | 60 |
| **Defocus range (µm)** | -2.4 to -1.0 |
| **Pixel size (Å)** | 0.5325 |
| **Symmetry imposed** | C 1 |
| **Movies (no.)** | 5291 |
| **Software for data collection** | EPU |
| **Initial “particle” images (no.)** | 4,844,663 |
| **Protein particle images (no.)** | 2,217,300 |
| **Final particle images (no.)** | 502,247 |
| **Map resolution unmasked (Å)** | 3.6 |
| **Map resolution masked (Å)** | 3.1 |
| **FSC threshold** | 0.143 |
| **Refinement** |  |
| Initial model used (PDB code) | 8TMU |
| Model resolution (Å) | 3.36 |
| FSC threshold | 0.5 |
| **Model composition** |  |
| Non-hydrogen atoms | 8073 |
| Protein residues | 1040 |
| **B factors (Å<sup>2</sup>)** |  |
| Protein | 86.65 |
| **R.m.s. deviations** |  |
| Bond lengths (Å) | 0.003 |
| Bond angles (°) | 0.509 |
| **Validation** |  |
| MolProbity score | 1.65 |
| Clashscore | 5.67 |
| Poor rotamers (%) | 0.57 |
| **Ramachandran plot** |  |
| Favored (%) | 95.01 |
| Allowed (%) | 4.99 |
| Disallowed (%) | 0.00 |
| **EMRinger score** | 2.82 |

**Table S4.** Data processing and refinement statistics for HLA-B*73:01 with KP1 and KIR2DL2.

|  |  |
| --- | --- |
| Wavelength | 1.0332 |
| Resolution range | 54.3 - 2.9 (3.004 - 2.9) |
| Space group | P 43 21 2 |
| Unit cell | 92.69 92.69 200.98 90 90 90 |
| Total reflections | 545044 (20136) |
| Unique reflections | 20180 (1973) |
| Multiplicity | 22.4 (16.8) |
| Completeness (%) | 99.97 (100.00) |
| Mean I/sigma(I) | 11.9 (0.9) |
| Wilson B-factor | 68.74 |
| R-merge | 0.199 (3.260) |
| R-meas | 0.203 (3.362) |
| R-pim | 0.043 (0.813) |
| CC1/2 | 1.00 (0.411) |
| Reflections used in refinement | 20177 (1973) |
| Reflections used for R-free | 999 (94) |
| R-work | 0.2225 (0.3146) |
| R-free | 0.2551 (0.3436) |
| Number of non-hydrogen atoms | 4764 |
| Macromolecules | 4645 |
| Ligands | 59 |
| Solvent | 60 |
| Protein residues | 588 |
| RMS(bonds) | 0.003 |
| RMS(angles) | 0.6 |
| Ramachandran favored (%) | 95.33 |
| Ramachandran allowed (%) | 4.67 |
| Ramachandran outliers (%) | 0 |
| Rotamer outliers (%) | 3.08 |
| Clashscore | 7.7 |
| Average B-factor | 70.86 |
| Macromolecules | 70.83 |
| Ligands | 80.78 |
| Solvent | 63.27 |
| Number of TLS groups | 15 |

**Table S5.**
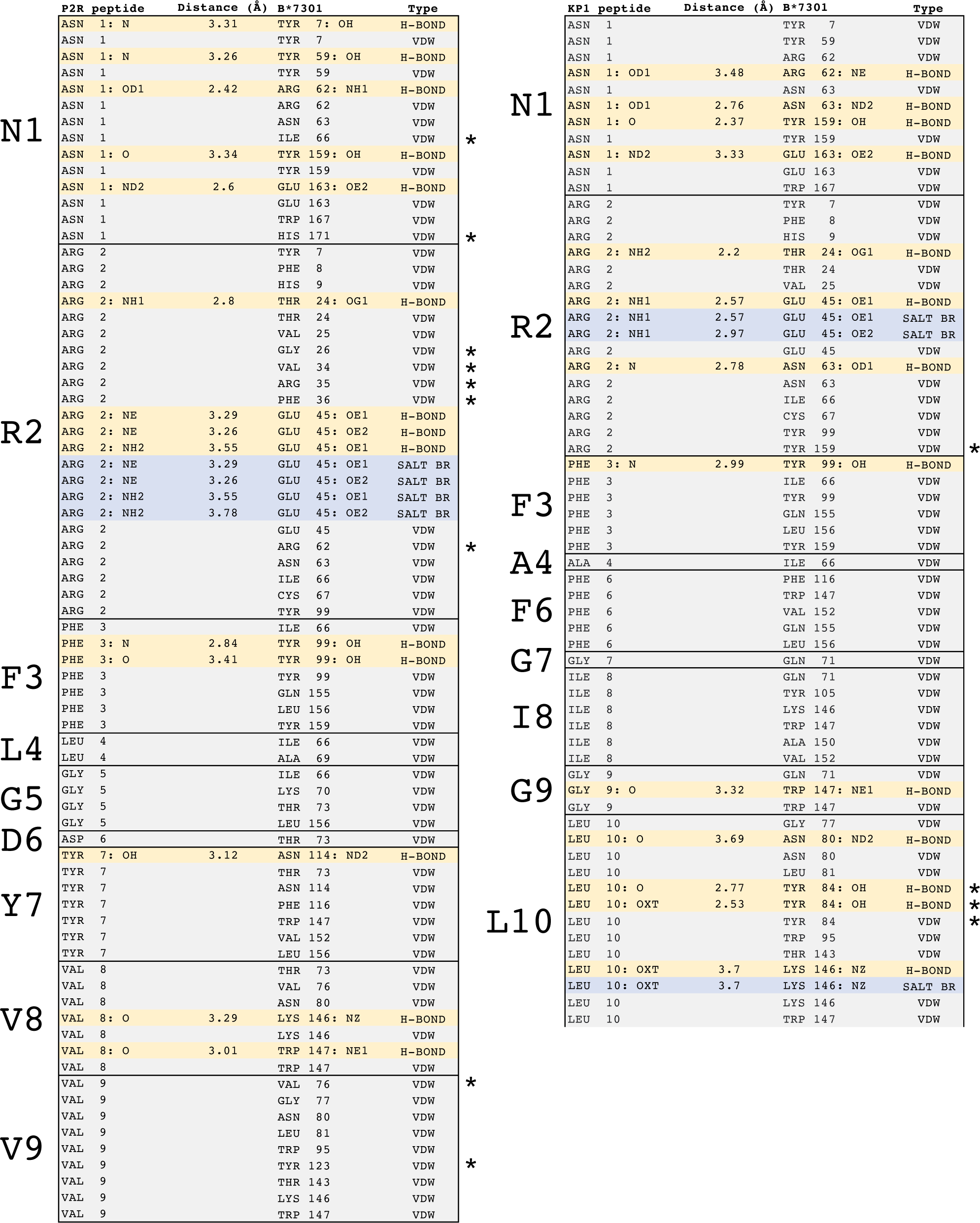
Peptide:B*7301 contact table.

